# Missingness Adapted Group Informed Clustered (MAGIC)-LASSO: A novel paradigm for prediction in data with widespread non-random missingness

**DOI:** 10.1101/2021.04.29.442057

**Authors:** Amanda Elswick Gentry, Robert M. Kirkpatrick, Roseann E. Peterson, B. Todd Webb

## Abstract

The availability of large-scale biobanks linking rich phenotypes and biological measures is a powerful opportunity for scientific discovery. However, real-world collections frequently have extensive non-random missingness. While missing data prediction is possible, performance is significantly impaired by block-wise missingness inherent to many biobanks. To address this, we developed Missingness Adapted Group-wise Informed Clustered (MAGIC)-LASSO which performs hierarchical clustering of variables based on missingness followed by sequential Group LASSO within clusters. Variables are pre-filtered for missingness and balance between training and target sets with final models built using stepwise inclusion of features ranked by completeness. This research has been conducted using the UK Biobank (n>500k) to predict unmeasured Alcohol Use Disorders Identification Test (AUDIT) scores. The phenotypic correlation between measured and predicted total score was 0.67 while genetic correlations between independent subjects was high >0.86, demonstrating the method has significant accuracy and utility.

## Introduction

Biobanks are large-scale, high-dimensional collections of biomedical information offering significant opportunities for scientific discovery, with many collections containing thousands of data points on tens of thousands of individuals. Many biobanks also collect biospecimens and perform genome-wide assessments of genetic variation and increasingly other omic measures such as gene expression, epigenetic modifications, and proteomics which allow comprehensive agnostic investigations of the relationships between complex human diseases and traits with genetic and environmental influences. These powerful resources are increasingly accessible to the larger scientific community, facilitating novel investigations and discovery. The breadth of phenotypes in biobanks represents an opportunity for machine learning (ML) approaches to further discover unexpected relationships complementing directed *a priori* hypothesis testing. However, the scale of biobanks also presents challenges including significant missing data, much of which is non-random.

There is a growing list of available biobanks for epidemiological discovery including the UK Biobank^1^ (UKB) which has enrolled over half a million UK residents, all of whom provided biological samples for genotyping. Volunteers in the UKB also provided access to their electronic health records, hospitalization records, biological samples, and answers to survey questions regarding diet, lifestyle habits, and mental health; phenotypic measures available to link with genetic measures totaling in the thousands. In the US, the National Institutes of Health is funding the All of Us^2^ biobank effort, which as of September 2022 has enrolled 372,380 of its goal of one million participants who will provide biological samples, genotypic data, electronic health records, and answers to several series of survey questions. Similarly, BioBank Japan^3^ has sampled over 200,000 participants with one of 47 common diseases and collected genetic information along with health records and other phenotypic information. Many additional biobanks are currently available to researchers and construction of new biobanks continues, motivated in part by the necessity of collecting large sample sizes to study the genetics of complex traits.

Structural characteristics of biobanks present challenges for data analysis. Many biobanks do not administer every test or survey to each participant, as budget considerations, for example, often dictate how many participants receive more costly testing, such as imaging. In order to mitigate dropout and participant fatigue, a subset of questionnaires may be sent to each participant; requests for participation in a particular survey may have been sent to a portion of subjects and only a subset of those were returned. Similarly, subsets of subjects may be chosen to participate in additional surveys according to previous responses, where the decision logic for these selections may not be clear or available to researchers. These practices, while pragmatic for cost and volunteer retention, may result in widespread, block-wise missingness across the full biobank, in which large subsets of the full sample have completely missing values for a portion of question categories. This missingness is non-random across the full set of available measures in the biobank such that no subset of subjects with complete information exists in the sample. **Figure 1** illustrates the pairwise correlation between missingness patterns across over 1,000 UKB variables that had the highest observation counts. Similarly, **Figure 2** illustrates the pairwise complete observation count across the same UKB variables, demonstrating that there is significant missing data and that these patterns of missingness are correlated across the dataset.

**Figure 1:**
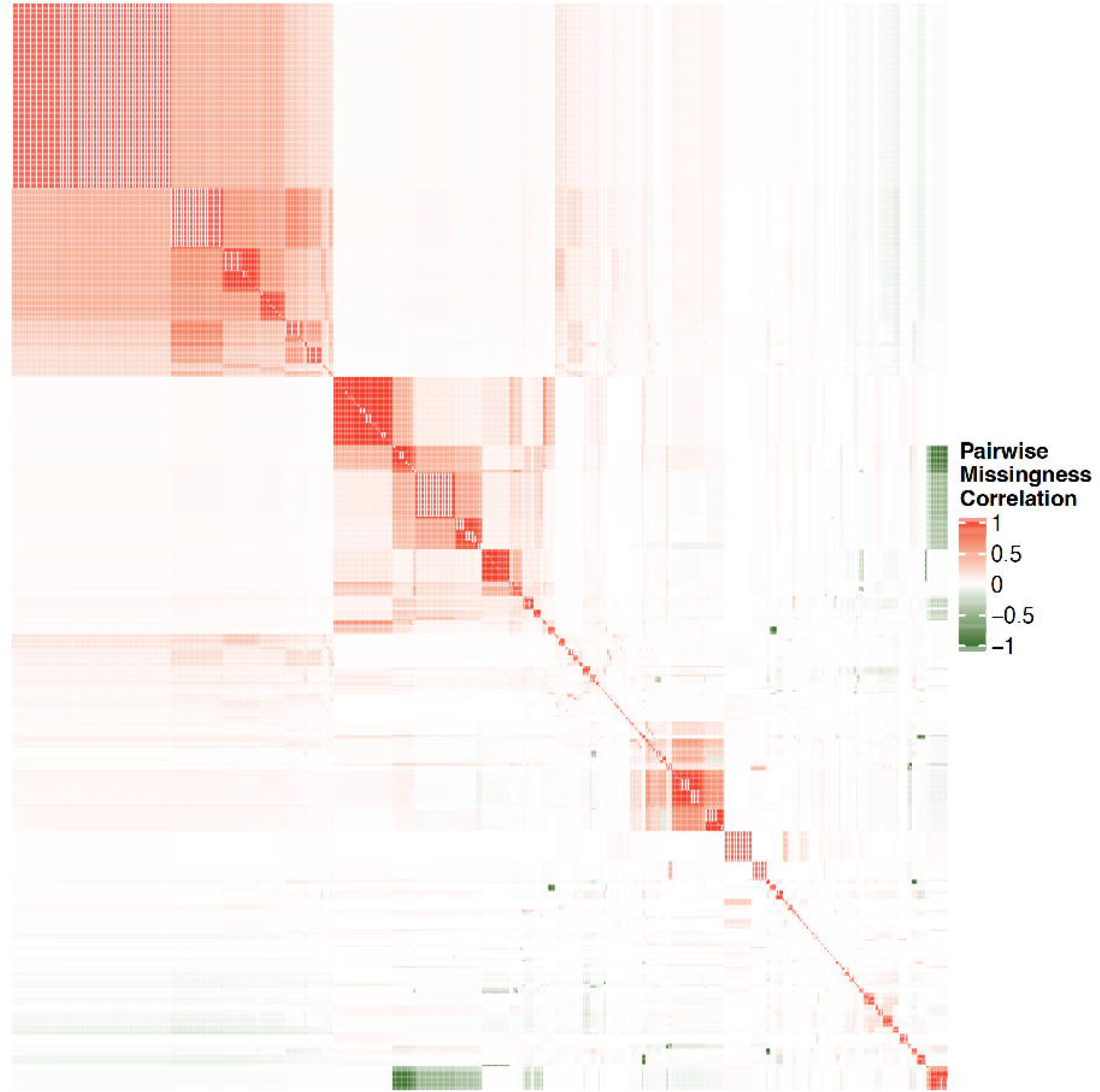
pairwise correlation between missingness patterns across over 1000 UKB variables with highest observation counts.

**Figure 2:**
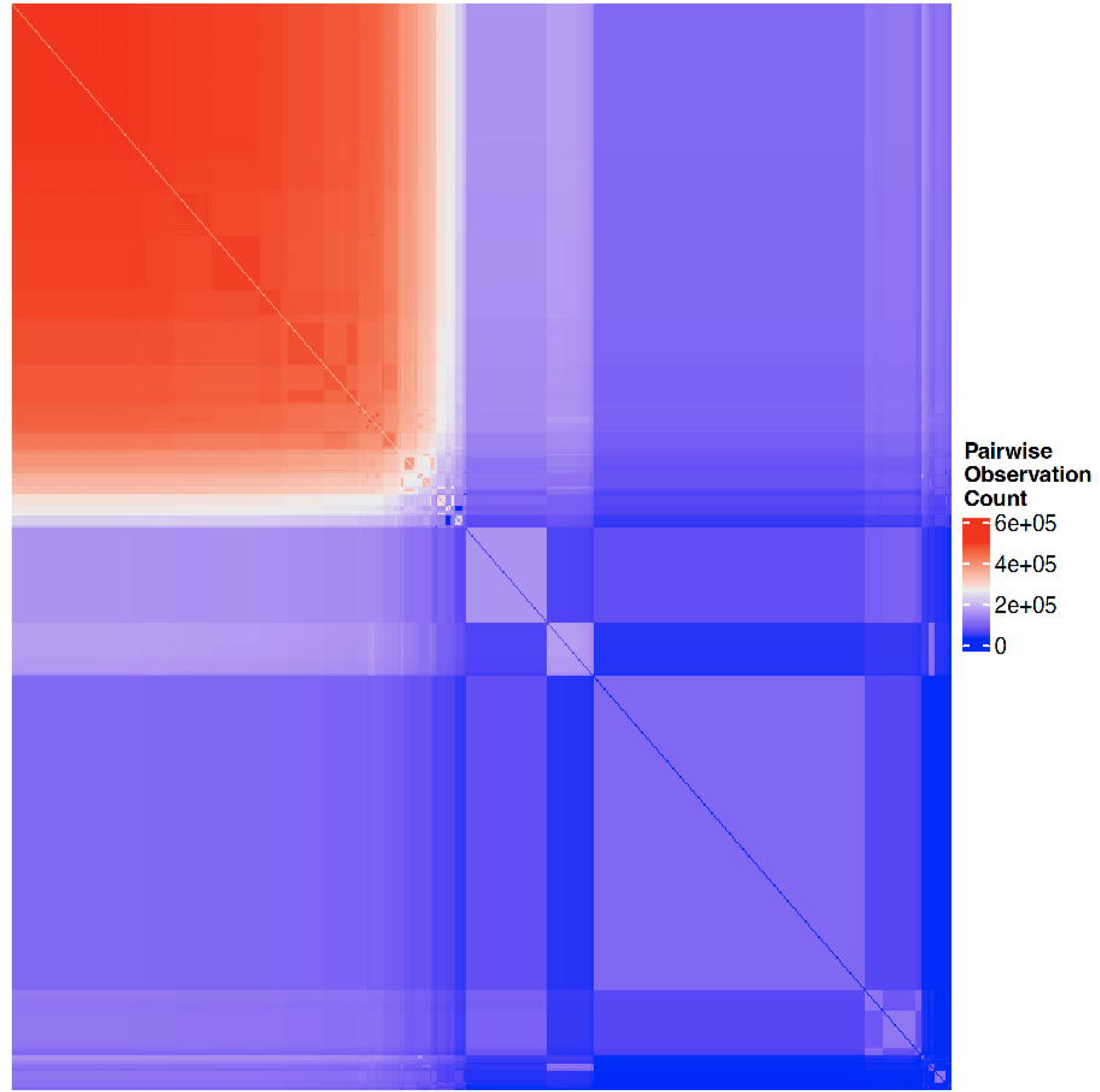
pairwise complete observation count across the same UKB variables shown in Figure 1

Missingness patterns can severely limit power for epidemiological and genetic analyses of any single trait. Traditionally, data missingness can be addressed through imputation where a missing-at-random structure can be reasonably assumed. Some commonly used approaches include *k* nearest neighbors^4^ or Multivariate Imputation by Chained Equations^5^ (MICE). These methods borrow information across the available data to infer missing points, but because biobank missingness is generally pervasive across all predictors and often decidedly non-random, these traditional imputation methods are not appropriate for estimating the missing values. Where imputation is inappropriate, row-wise deletion is sometimes employed to drop subjects who have missing observations. However, with even moderate levels of random and block-wise missingness, this sort of deletion can render the dataset many orders of magnitude smaller.

Given the phenotypic depth of biobanks, when traditional imputation may not be employed, there is an opportunity to apply ML methods to leverage the existing data to predict missing values. Utilization of ML, or “data mining,” as it is often called, has continued to rise across many applications, including genetics. For example, the the PsychENCODE project^6^ employed deep learning techniques to predict functional ramifications in the brain of genome-wide association study (GWAS) hits associated with psychiatric disorders. Advances in technology and cloud resources continue to ease the computational burden of applying ML methods to high-dimensional genomic data and offer the opportunity for rapid and novel investigations. While these advances have sparked wide-spread interest, many ML methods themselves are not novel, but rather are based in statistical techniques long established theoretically and proven empirically^7^.

Many ML procedures may be useful for predicting missingness in an outcome where there is only moderate and random missingness across the full set of predictors. When missingness is pervasive and non-random, as observed in many biobanks, traditional implementation of most data mining techniques also resorts to row-wise deletion of subjects with any missing predictor values. A subset of ML approaches have been adapted to account for some level of predictor missingness and applied to missing variable imputation. MI-LASSO^8^, for example, integrates Multiple Imputation (MI) of missing predictors with the Least Absolute Shrinkage and Selection Operator (LASSO) for a hybrid approach applicable where missingness may be assumed to be random. PhenIMP^9^ and extensions^10^ use related phenotypes to impute a difficult to collect phenotype in order to boost power. While PhenIMP can impute using only summary information from other phenotypes, it relies on distributional assumptions which make the approach impractical where many phenotypes are categorical and do not conform to such assumptions.

Similarly, the PHENIX^11^ method was designed to impute missing variables in a Bayesian framework in the presence of other informative data but also requires distributional assumptions and does not drop non-informative input measures, thereby prohibiting variable selection. Other approaches developed by Yuan et al.^12^ and expanded upon by Xiang et al.^13^ specifically address block-wise missingness structures with a focus on imputing entire blocks of missing data, specifically where neuroimaging data is present. While innovative and effective for applications involving a small number of well-defined blocks of data, this method is not applicable to the structure of large-scale data wherein the block-wise missingness patterns are highly inconsistent across subjects and the number of blocks is large.

Given the variety of available ML approaches and characteristics of biobanks, there is significant need for an ML solution for imputing missing variables which collectively (1) is capable of including categorical and/or non-normally distributed predictors, (2) produces interpretable models, (3) incorporates penalization or variable selection such that it could be generalizable, and most importantly, (4) is applicable and robust in the presence of non-random, block-wise missingness. While many traditional ML methods could satisfy the first three interests, most are intolerant to missingness in the predictors, precluding ‘out-of-the-box’ application of available methods.

Through missing data simulations, we demonstrate the rate at which missing-at-random and block-wise missing data reduce the number of complete cases in a dataset to zero, thereby motivating the necessity of novel ML implementations to handle missing data in biobank scale collections. We further simulate datasets with a variety of random and block-wise missingness structures and apply our proposed ML innovation to test its performance empirically. As a proof of principle, we selected the UKB to serve as an example real-world application of our proposed ML innovation. The data freeze (UKB Application 30782, approval date September 3, 2018, using data baskets created September 28, 2019 and May 20, 2019) contained 9,613 phenotypes on 502,536 subjects. We chose the Alcohol Use Disorders Identification Test (AUDIT) survey from UKB as our target outcome; it was ascertained as part of the mental health battery of questionnaires and was directly measured in 157,162 (31.2%) participants. The AUDIT is a ten-item, self-administered screening instrument for alcohol problems containing three questions surveying consumption and seven items surveying problems related to alcohol which comprise the AUDIT-C and -P subscales^14,15^. Here, we describe an innovative ML approach and demonstrate its usefulness in leveraging thousands of measured variables in order to predict an unknown, unmeasured variable and show how this predicted outcome boosts power for subsequent analyses including GWAS, cross-trait genetic correlation, and other statistical studies.

## Methods

### Data: UK Biobank

The UK Biobank is a large-scale biomedical database and research resource containing genetic, lifestyle, and health information from half a million UK participants. UKB’s database, which includes blood samples, heart and brain scans and genetic data of the volunteer participants, is globally accessible to approved researchers who are undertaking health-related research that is in the public interest. Participants were aged between 40-69 years and recruited in 2006-2010 from across the UK. This research has been conducted using the UK Biobank Resource application number 30782.

### Missing Data Simulations

To demonstrate the relationship between complete case count and both random and block-wise missingness, we simulated an indicator matrix with 200 variables on 10,000 subjects and imposed random missingness from 1-50% of the data and block-wise missingness in 0, 5, 10, 15, and 20 blocks ranging in size from 5-15 columns and 100-500 (in increments of 5) rows. This missingness was randomly imposed over 100 iterations so that median complete case count could be calculated across the missingness parameters.

### MAGIC-LASSO

We developed an adaptation of the Group Least Absolute Shrinkage and Selection Operator^16^ (Group-LASSO) machine learning method for penalized regression to address the shortcomings of existing, software-implemented ML methods for predicting variables in the presence of non-random, block-wise missingness named the Missingness Adapted Group Informed Clustered (MAGIC)-LASSO. The MAGIC-LASSO represents an innovative implementation of established ML methods and is therefore intended to serve as a new *application* of existing algorithms. While other choices of ML approach may have been appropriate, for this first proof of principle project, we chose to extend and adapt the Group LASSO because it employs a straightforward and easily interpretable regression-based solution and it is particularly suited for penalization of categorical predictors, of which there are many in the UKB.

Details regarding the theory, formula, and derivation of the LASSO^17^ and one of its extensions, the Group-LASSO^16^ are published elsewhere and are summarized in **Supplementary Material, Section 1**. Our MAGIC-LASSO approach utilizes the conventional Group-LASSO fitting algorithm, but applies it in an innovative, iterative manner in order to overcome the challenges of the block-wise missingness design.

### MAGIC-LASSO overview

In brief, the MAGIC-LASSO procedure involves (1) characterizing missingness, (2) filtering variables for general missingness and for balance across training and target sets, (3) variable clustering based on missingness, (4) iterative Group-LASSO and variable selection within clusters, and (5) cross-cluster model building with variables prioritized by informativeness. **Figure 3** describes the flow logic of the MAGIC-LASSO.

**Figure 3:**
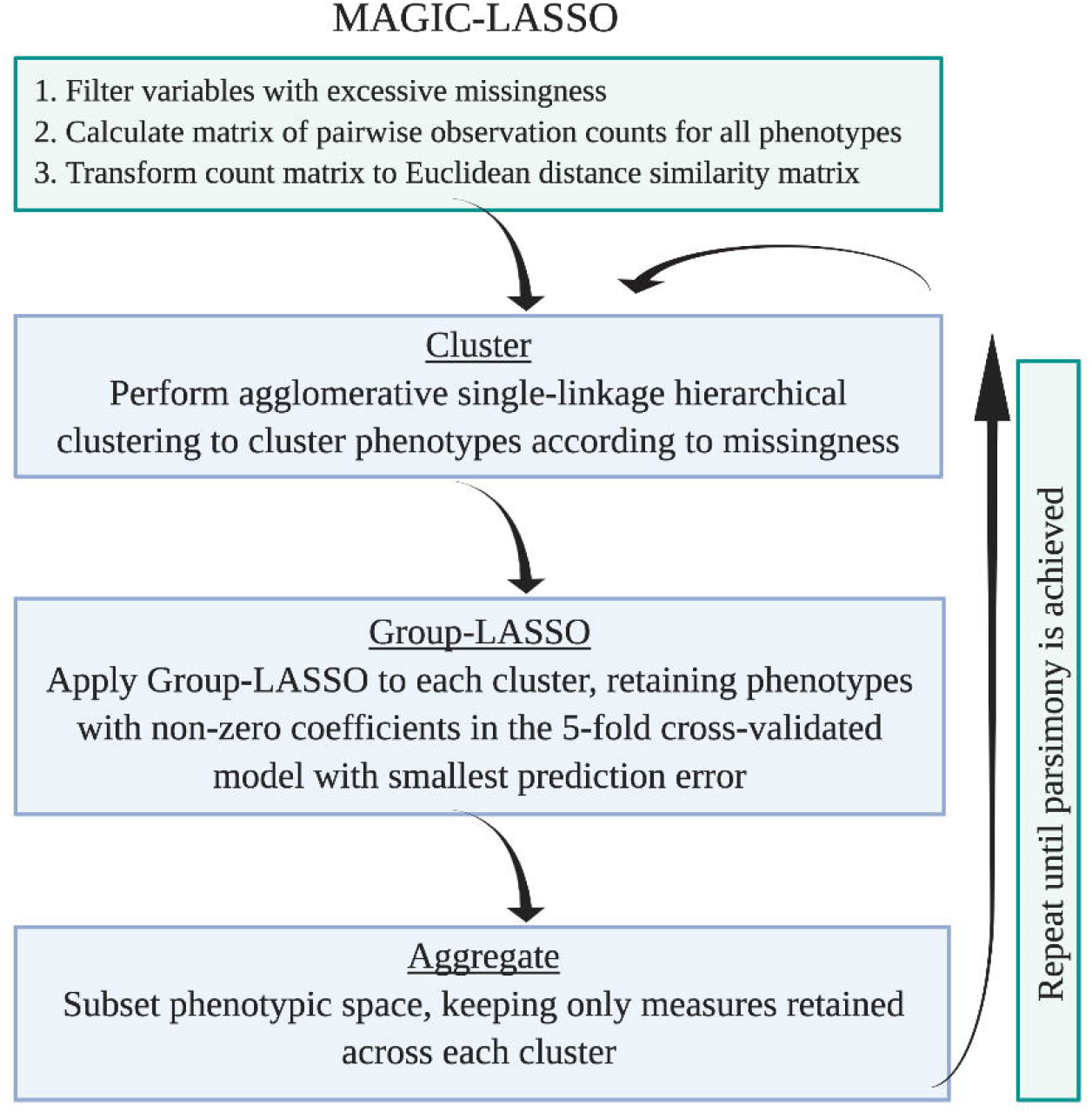
Flow logic of the MAGIC-LASSO procedure. The MAGIC-LASSO procedure begins with filtering, followed by clustering, then iterative Group-LASSO application until parsimony is achieved.

### Characterizing missingness and general filtering

This first step is to create a subset of variables suitable for downstream investigation. This includes removing potential predictor variables that are (a) excessively sparse (> 80% missingness), (b) categorical with excess, sparse levels such as ICD codes in a collapsed matrix format, (c) unstructured where the format is inappropriate for modeling, such as free text, date values, or array variables, or (d) invariant. After initial filtering, we identify and remove variables for which missingness patterns were highly skewed between prediction and training sets for the outcome of interest. Due to block-wise missingness, there may be variables which pass the first filtering step but are not informative in the target dataset. In other words, where data completeness is highly correlated with the variable of interest. This is not to be confused with correlation among the phenotypic measures themselves, which is generally not of concern since the LASSO procedure is more capable of handling many measures with varying degrees of collinearity than traditional linear regression^7^.

### General background on ML training and test sets

This filtering step relies on the identification of a so-called measured set, also referred to here as a training set, the subset of subjects with the primary outcome measured, and an unmeasured set, or a prediction set, the remaining subjects for whom the outcome of interest was unmeasured and for whom the ML procedure will predict the missing variable. **Figure 4** illustrates an example of how a dataset may be subdivided into these measured and unmeasured sets.

**Figure 4:**
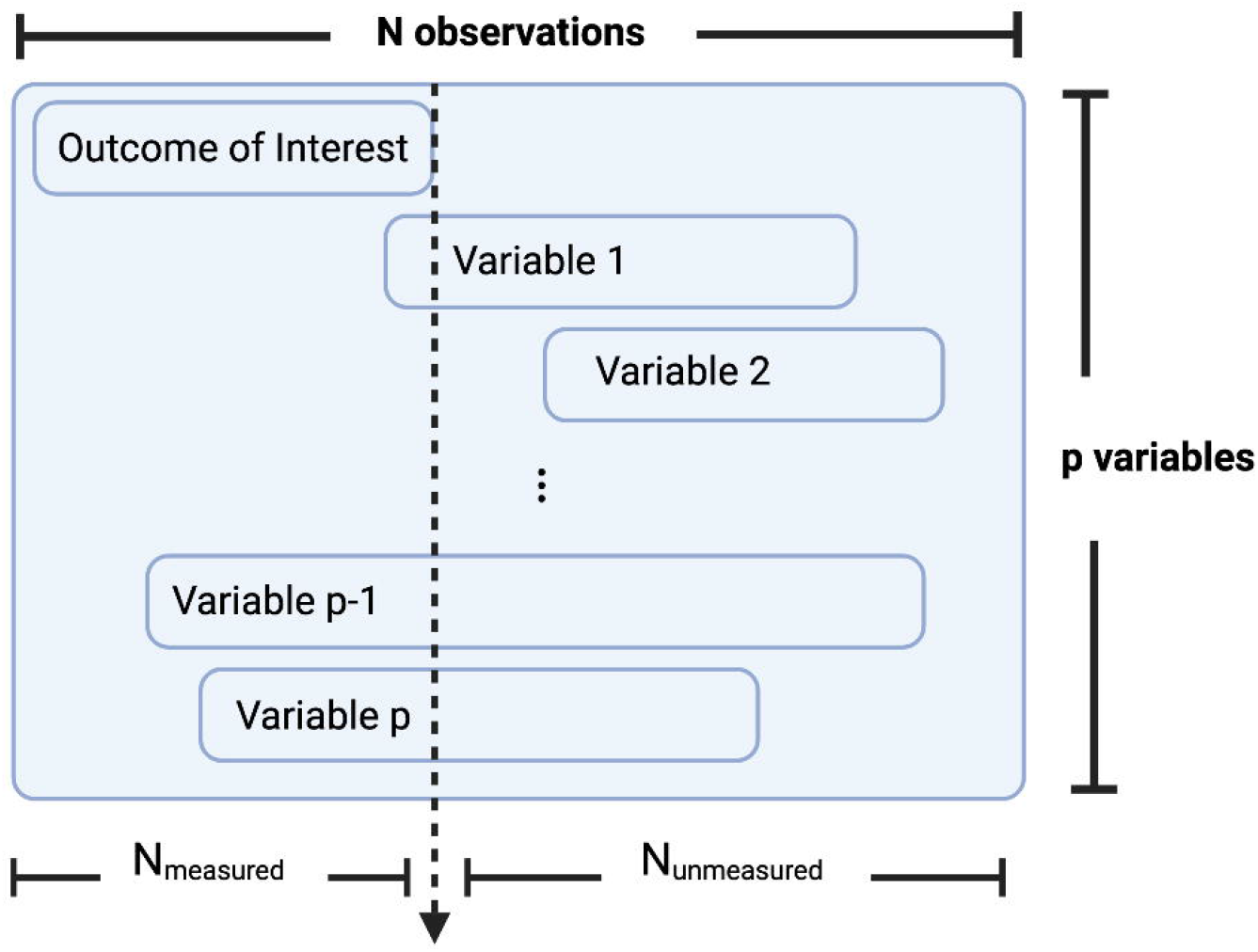
Conceptualization of how a dataset may be subdivided into a measured and unmeasured set. Where *N* represents the full sample size, *N*_*Unmeasured*_ and *N*_*Measured*_ represent the subsets of subjects on whom the outcome of interest is either missing or measured, respectively. Then the amount of overlap in observation may be quantified for each of p additional variables.

### Balancing

When training an ML model to predict unmeasured variables, the learning occurs on the subset of data for which complete observations are available, i.e., the measured, or training set and is then implemented in the unmeasured, or the prediction set. The algorithm learns how to predict unobserved data by modeling patterns that exist in observed data. Where certain variables are largely measured in conjunction with the primary outcome of interest in the training set but are largely unmeasured in the prediction set, an ML algorithm which relies on these variables for prediction will perform poorly, since the inputs will be largely missing.

For a given experiment, partition the total number of observations into those in the measured and unmeasured sets *N*_*measured*_ + *N*_*unmeasured*_ = *N*_*total*_ and for each additional variable *k*, quantify *n*_*k,measured*_ and *n*_*k,unmeasured*_ the number of observations present in *N*_*measured*_ and *N*_*unmeasured*_, respectively. Then calculate a filtering parameter:

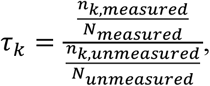

where *τ*_*k*_ represents the ratio of the proportion of observations present in the measured set to the proportion of observations present in the unmeasured set, for variable *k*. Plotting is helpful to empirically determine a useful cutoff for *τ* = *t*.

### Clustering

The Group-LASSO, like many ML procedures, cannot accommodate missing data and relies on row-wise deletion of observations where one or more variables are missing. One strategy to mitigate reduction in sample size from requiring complete information across all covariates is to segregate the variables into blocks according to patterns of missingness and apply the ML procedure within that subset of measures. In the MAGIC clustering step, variables are grouped to minimize missingness while maximizing sample sizes in order to optimize downstream within-cluster prediction performance. First, pairwise observation counts for every pair of phenotypic variables are calculated. Using this pairwise count matrix, we calculated the Euclidean distance of these measures to feed into an average-linkage agglomerative hierarchical clustering procedure to discover the inherent groupings of variables based on missingness. This clustering procedure begins with each variable in its own cluster and proceeds by combining two clusters for each step until all variables reside in a single cluster. The clustered tree may be cut at some point to obtain the clustering assignments. Exact height for cutting is determined empirically by examining mean observation count per variable in the cluster, number of variables in the cluster, and the number of complete cases for that subset of measures.

### Iterative Group-LASSO

The cut tree provides groups of variables within which the complete data observation count is maximized. Limiting each cluster to only the complete data therefore, the Group-LASSO is applied to each cluster individually. Each model utilizes *k*-fold cross-validation to choose the penalty term with minimal prediction error and variables in each model were retained if they achieved non-zero effect estimates, where *k* is chosen to be small and approaching *n*, with consideration of computation resources. After applying the Group-LASSO to each cluster, variables retained by each model are aggregated across clusters, as illustrated in **Supplementary Figure 1**. The set of aggregated, retained variables is then re-clustered using the same hierarchical clustering procedure and the Group-LASSO applied to each cluster. With each successive iteration of the clustering and Group-LASSO application, the predictor space shrinks as the measures most predictive of the outcome are retained across iterations and the less-informative measures are dropped.

**Figure 3** illustrates the flow of the algorithm, which continues until moderate parsimony is achieved. We recommend stopping when (1) repeated iterations no longer shrink the predictor space or (2) the data set with the retained predictors has a reasonable number of complete cases. The number of complete cases needed to fit a final iteration of the model will vary depending on the total number of predictors, the total sample size, and the variation in the data, but the Group-LASSO is capable of effectively fitting a model even where *p*, the number of variables, is larger than *n*, the number of samples.

### Cross cluster model building

Once the iterative procedure is halted, with *p* remaining variables, the data is split into independent training and test sets and the Group-LASSO is fit up to *(p* − 1) times using a stepwise procedure which orders the variables according to missingness. Beginning with the variable with least missingness and adding an additional variable each round, the Group-LASSO model is fit to the training set complete data on those variables and *k*-fold cross-validation is used to determine the phenotypic correlation between the observed and predicted outcome. With each step, an additional variable is added and the Group-Lasso fit, and the procedure continues until every set has been fit, or there are no longer any complete cases in the successive set. The set of variables producing the most predictive model in the hold-out test set is chosen as the final model.

#### Algorithm

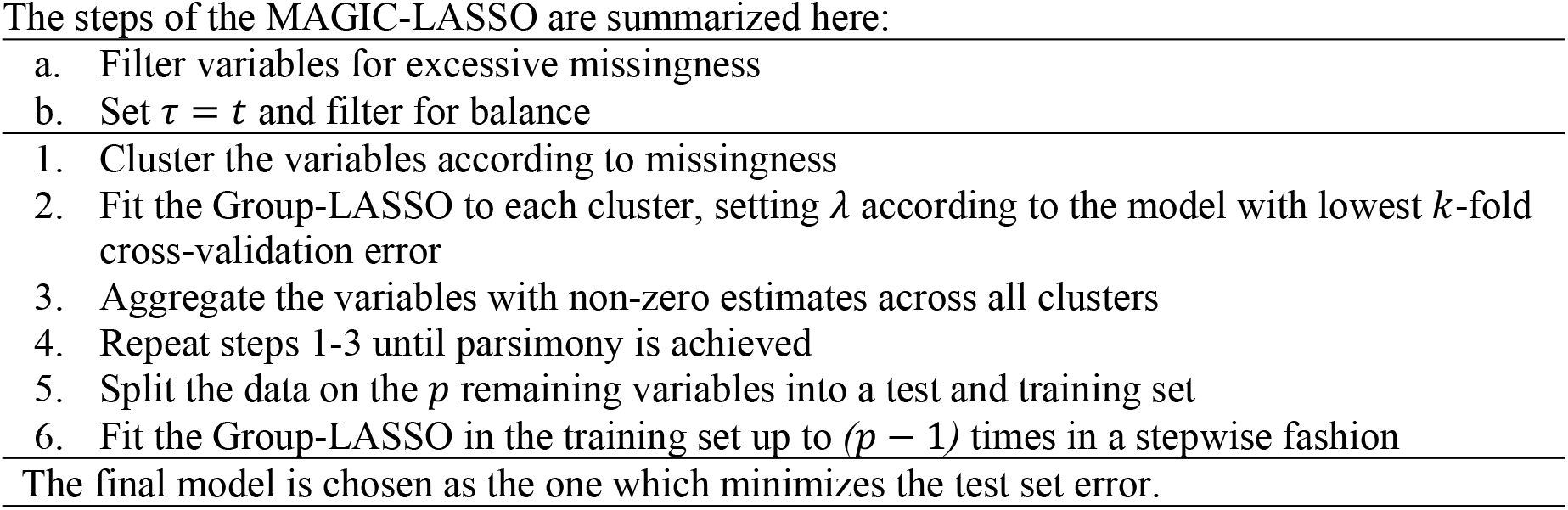

### Test and training sets

Although each iteration of the Group-LASSO application is fit using *k*-fold cross-validation, it is optimal to further utilize a hold-out test set during the construction of the final Group-LASSO model in order to rigorously assess performance. The proportion of the data assigned to the hold-out test set depends on the size of the data set itself, although a hold-out set containing 10-30% of the data is typical^7^. Phenotypic predictive performance is assessed by plotting observed versus predicted observations in the test set and reporting the correlation between the observed and predicted sets.

### MAGIC-LASSO simulations

To test the prediction performance of the MAGIC-LASSO approach empirically, we applied the modeling procedure to data simulated according to the following steps:

1. Beginning with 354 UKB variables selected and filtered for the AUDIT application as described above, calculated ***σ***_**0**_, the 354 *x* 354 variance/covariance matrix and ***μ***, the vector of means
2. Filtered measures with excessive variation, resulting in 299 remaining measures
3. Applied the *nearPD()* from the Matrix R package to find the closest positive-definite variance/covariance matrix, ***σ***
4. Generated multivariate normal data from ***σ*** and ***μ***

Data was generated using sample sizes of *N*=10,000 or 50,000 and *P*=200 or 250 phenotypes. Half of the predictors were selected to be categorical with either 4 or 6 categories. The outcome measure was generated via a regression equation using either 5 or 10 phenotypes and error following a *N*(0,2) distribution. We utilized tree cut points of either (6, 7, 8, 9, 10) or (3, 4, 5, 6, 7, 8, 9, 10, 11, 12). In order to fully capture the complex, correlated nature of the blockwise missingness structure inherent to biobank data, we further imposed on the simulated data set the missingness structure observed from a random subset of the true data, in addition to imposing various levels of random missingness of either (500, 1000, 1500) or (2500, 5000, 7500) observations. Performance was evaluated by comparing the correlation and mean squared error (MSE) between the observed and predicted outcomes for both the set of missing outcomes and for the full set of outcomes. Further details of the simulated data may be found in the **Supplementary Material, Section 2**.

### GWAS

For our real-world data application, to assess how well the predicted AUDIT outcome captures the underlying genetic factors influencing AUDIT, we calculate the heritability of observed and predicted AUDIT as well as the genetic correlations (*r*_*g*_) between the observed and predicted outcomes. To this end, we conducted GWAS (bgenie^18^ version 1.3) of the AUDIT-Total, AUDIT-C, and AUDIT-P scores in the measured and the combined measured-plus-unmeasured sets. We utilized common procedures for pre-GWAS filtering, including excluding markers with MAF < 0.5%, INFO < 0.8, and HWE p-value < 10^−6^. All association analyses included age, sex, and the first 20 ancestry principal components as covariates. The independent European subjects sample size for the GWAS was 359,980 with 117,559 and 242,421 subjects in the AUDIT measured and unmeasured sets, respectively.

### Heritability and Genetic Correlation

GCTA^19^ (version 1.93.2) was used to calculate heritabilities and the *r*_*g*_ between observed and predicted AUDIT, but only within the set of participants on whom AUDIT was directly measured since the GCTA only allows *r*_*g*_ to be calculated across the same set of observations. We also utilized LDSC^20,21^ (version 1.0.1) to estimate heritabilities and *r*_*g*_ between observed and predicted scores, both within the measured set and between the measured and unmeasured sets. Using LDSC, *r*_*g*_ can be estimated in either independent or overlapping samples by leveraging a reference set of genetic correlations (linkage disequilibrium) and GWAS summary level test statistics.

### Software

Data management and application of the MAGIC-LASSO was conducted in R^22^ (v3.5.2) using packages *Matrix*^23^ (v1.2.17), *fastDummies*^24^ (v1.5.0), and *grpreg*^25,26^ (v3.2.1). Clustering was conducted using *hclust* UPGMA method in base R and the Group-LASSO was fit using the *cv*.*grpreg* function in the *grpreg* package. R scripts describing the MAGIC-LASSO implementation are available at *github*.*com/AEGentry/MAGIC_LASSO*.

## Results

### Simulation results

The results of the missingness simulations indicated that regardless of the level of blockwise missingness, once random missingness across the dataset reached 5%, the median number of complete cases fell to 0. In fact, the number of missingness blocks affected the complete case count less than the overall proportion of random missingness, as illustrated in **Figure 4**. The full table of these simulation results may be found in **Supplementary Table 1**.

The full results of the MAGIC-LASSO simulations may be found in **Supplementary Tables 2-7**. In summary, for models with some complete cases present, the MAGIC-LASSO performed better than the regular Group-LASSO, achieving overall lower MSE and higher correlation between observed and predicted outcomes in the complete model in all scenarios. In cases with zero complete cases present, performance between the regular Group-LASSO and MAGIC-LASSO could not be compared because the Group-LASSO is incompatible with missing data in the predictors. In the zero complete cases scenarios, correlation between observed and predicted outcomes ranged between 0.667 and 0.760. While prediction performance was good across scenarios, we observed higher overall correlations and correspondingly lower MSE for scenarios 1, 3, 4, and 6, in which the true beta values were smaller in magnitude than those in scenarios 2 and 5. The choice of optimal cut point for the clustering trees varied across scenarios, indicating that no one cut point is objectively optimal in all scenarios. Furthermore, in most scenarios, predictive performance between cut points did not vary widely.

### Application to real world biobank data

As a proof of principle, we applied MAGIC-LASSO to predict AUDIT in the UKB. We predicted the subscales (AUDIT-P and AUDIT-C) individually, as well as the total AUDIT, calculated as the sum of the two subscales. The median total AUDIT score was 4 while the median for the consumption (AUDIT-C) and problem (AUDIT-P) subscales was 4 and 0, respectively.

To construct a set of variables to be used in the predictive algorithm, we filtered the full set of 9,603 available variables (not including the AUDIT measures) to remove measures (a) with fewer than 100,000 observations, (b) which were ICD codes, (c) which were unstructured, (d) which were invariant, or (e) which were repeated and measured at later longitudinal timepoints, such that only baseline measures were retained. **Supplementary Figure 2** shows the sample sizes remaining after each filtering step. After these filtering steps, 631 curated, so-called top-level (i.e., baseline) variables remained. Further filtering for balance between the measured and unmeasured sets removed 277 more variables for which missingness patterns were highly skewed between prediction and training sets. Measures with a ratio of missingness in the predicted versus the training set of *t* ≤ 0.7 were filtered out, leaving 354 variables.

Clustering the post filtered variable set resulted in an initial 12 clusters (**Supplementary Table 8**). One cluster of 5 variables was dropped because there were no complete cases in the cluster. Using 5-fold cross-validation, the first application of the Group-LASSO resulted in an aggregated total of 99, 106, and 123 variables retained across all clusters for the AUDIT-Total, AUDIT-C, and AUDIT-P, respectively. In the second iteration, variables were grouped in 5, 6, and 4 clusters for AUDIT-Total, AUDIT-C, and AUDIT-P respectively and applying the Group-LASSO to each cluster resulted in an aggregate of 65, 80, and 54 variables retained across the clusters for AUDIT-Total, AUDIT-C, and AUDIT-P, respectively.

Having reduced the phenotypic space by nearly a quarter for each score, the iterative Group-LASSO process halted. We then ordered the variables in each set according to missingness and applied the Group-LASSO procedure to the set of variables constructed in a forward stepwise manner, beginning with the variable with least missingness. **Supplementary Table 9** shows the number of subjects with complete data, with the addition of each variable, including the breakdown of complete cases in the measured and unmeasured sets, as well as the correlation from a predictive model constructed using each successive set of variables. The phenotypic correlations and the ratio of proportion of complete cases from the measured and unmeasured sets are shown for each outcome in **Supplementary Figure 3**. The stepwise procedure showed final models with 30, 18, and 20 input variables resulted in the best prediction for AUDIT Total, Consumption, and Problems, respectively. The final models resulted in 27, 18, and 14 non-zero coefficient estimates and test-set phenotypic correlations of 0.64, 0.71, and 0.48 for Total, Consumption, and Problems respectively.

Using the full measured sets for which both observed and predicted AUDIT scores were available, the phenotypic correlations were 0.65, 0.70, and 0.46 for Total, Consumption, and Problems, respectively. These estimates are very similar to those from the final models using only the test set which demonstrates the approach produces generalizable estimates. **Figure 6** shows the density curves of observed and predicted (both measured and unmeasured) for all three scores as well as the density curves of the prediction residuals.

**Figure 5:**
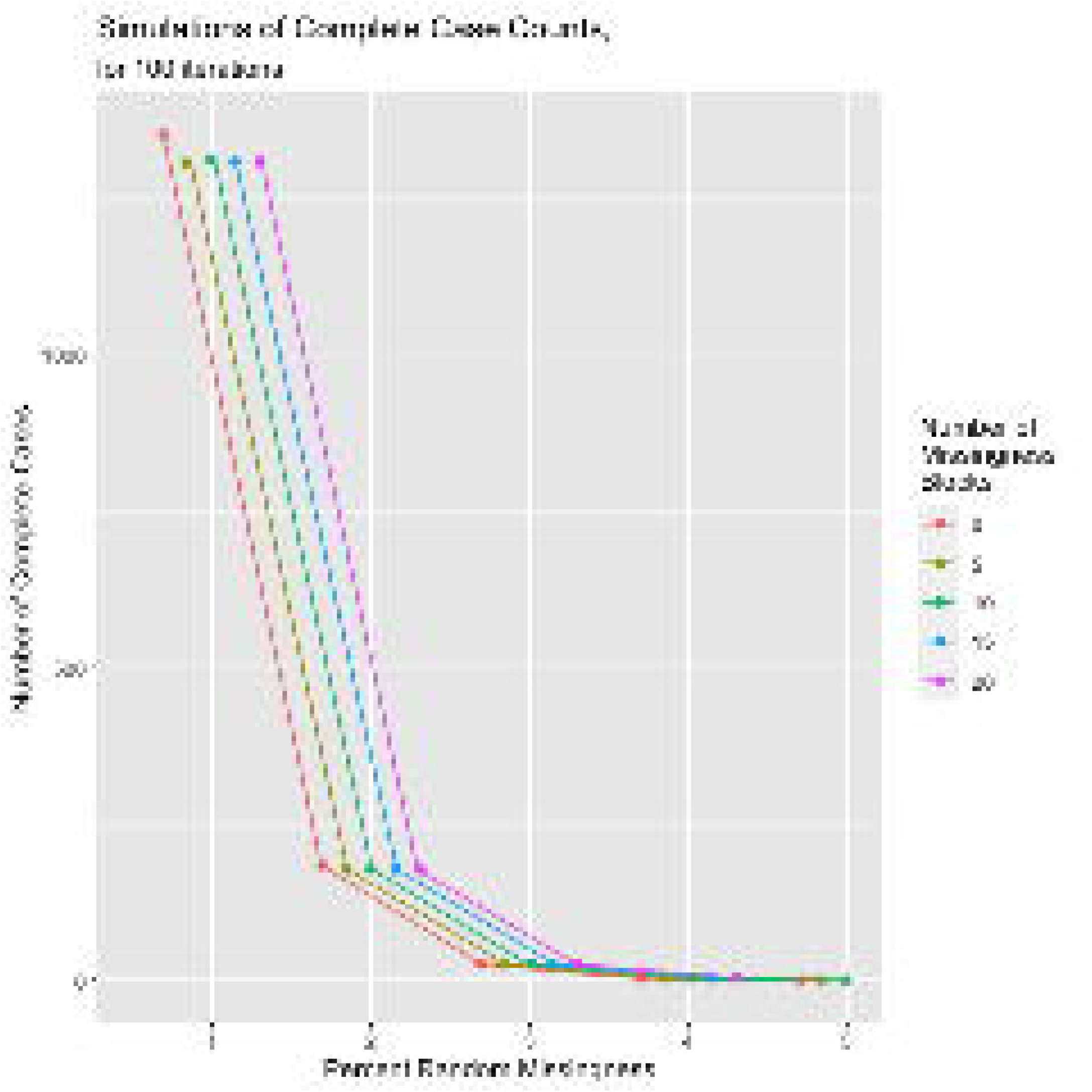
Number of complete cases as a function of number of missingness blocks and overall random missingness across the dataset.

**Figure 6:**
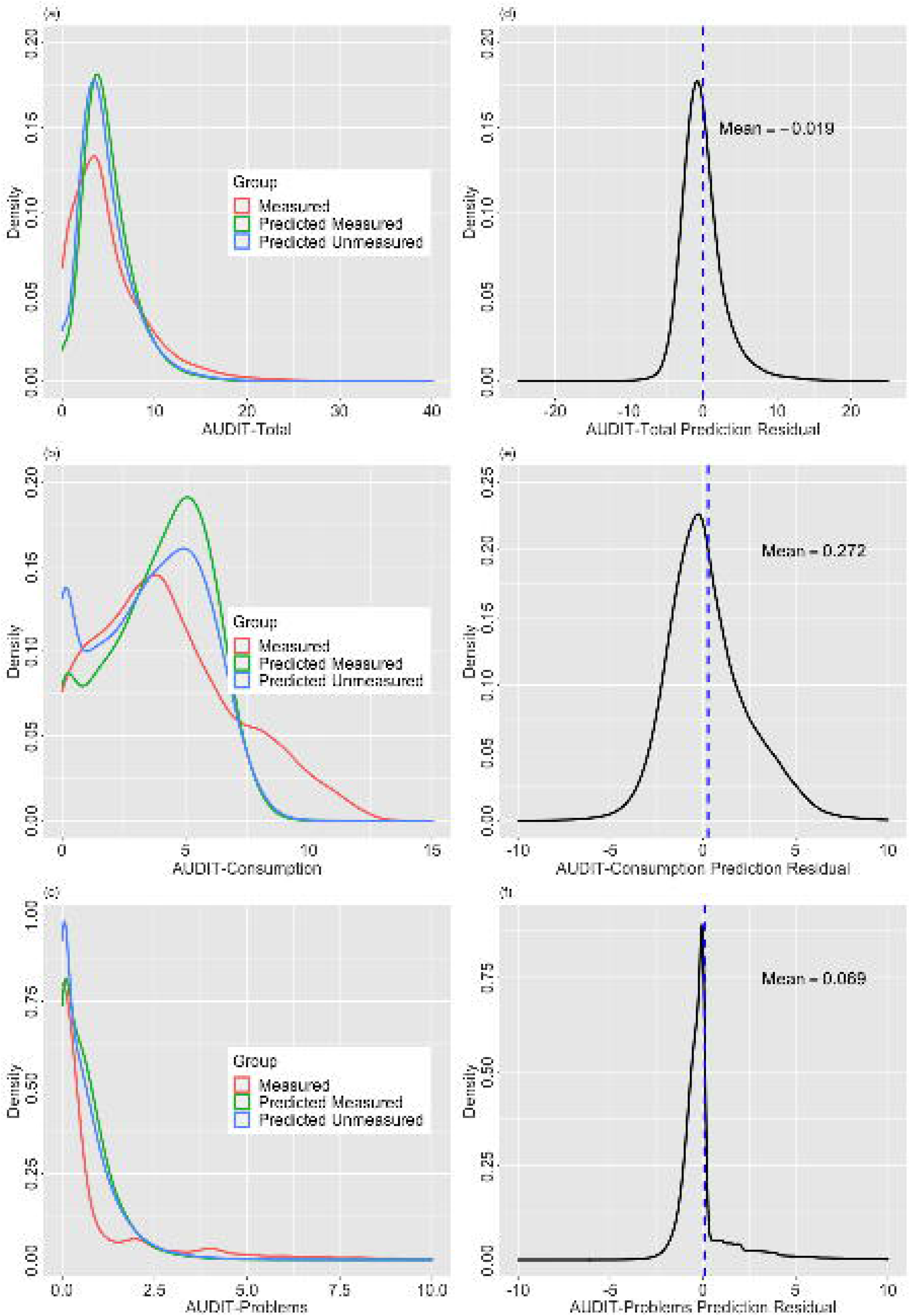
Densities curves showing observed and predicted outcomes and prediction residuals. **(Left)** Density curves of the observed and predicted scores; outcomes in the observed and predicted in the measured and unmeasured sets plotted for (a) AUDIT-Total, (b) AUDIT-C, and (c) AUDIT-C. **(Right)** Residual densities for AUDIT prediction; density curves with means noted showing the distribution of the prediction residuals for (d) AUDIT-Total, (e) AUDIT-Consumption, and (f) AUDIT-Problems.

One significant advantage to evaluating ML methods including MAGIC-LASSO in biobanks such as UKB is the availability of genetic information on all subjects and methods to estimate SNP-based heritabilities (*h*^2^) and genetic correlations (*r*_*g*_). To explore the accuracy of the predicted phenotypes and evaluate their utility in downstream genetic studies, we estimated *h*^2^ in each set and *r*_*g*_ between (a) observed and predicted in the subjects with measured AUDIT and (b) predicted AUDIT in subjects with and without direct measurement. We note that the last sets are completely independent with no information being shared in the model building step except for missingness balance.

### Heritability

Within-subject GCTA based *h*^2^ for AUDIT-T in men and women showed similar estimates when directly measured as when predicted by MAGIC-LASSO (range 0.089 – 0.139) (**Supplementary Table 10)** and was similar to LDSC based estimates (range 0.047 – 0.087) (**Table 1(a)**) derived from GWAS summary statistics. Of note, the estimated heritabilities of the predicted score are close to those of the observed score in both the measured and unmeasured sets.

**Table 1:** LDSC estimated heritabilities and genetic correlations. Block (a), heritabilities (*h*^2^) for observed and predicted AUDIT; p-values estimated from z-scores calculated using h2 and se estimates. Block (b), genetic correlations (*r*_*g*_) between the observed and unobserved AUDIT in the measured and unmeasured sets.

|  | **(a)** |  |  | **(b)** |  |  |
| --- | --- | --- | --- | --- | --- | --- |
| **Set** | **Outcome** | **$h^2$ (SE)** | **p-value** | **Comparison** | **$r_g$ (SE)** | **p-value** |
| **AUDIT-Total** |  |  |  |  |  |  |
| Measured | Observed | 0.0811 (0.006) | 1.25E-41 | Unmeasured<br>Predicted | 0.9746<br>(0.0354) | 5.40E-167 |
| Unmeasured | Predicted | 0.0727 (0.0036) | 1.10E-90 | Measured<br>Predicted | 0.9746<br>(0.0333) | 2.30E-188 |
| Measured | Predicted | 0.08433 (0.0056) | 3.02E-51 | Measured<br>Observed | 0.9191<br>(0.0181) | <1.0E-200 |
| **AUDIT-Consumption** |  |  |  |  |  |  |
| Measured | Observed | 0.0869 (0.0061) | 4.75E-46 | Unmeasured<br>Predicted | 0.8695<br>(0.0353) | 3.30E-134 |
| Unmeasured | Predicted | 0.0759 (0.0042) | 5.35E-73 | Measured<br>Predicted | 0.9712<br>(0.0349) | 1.40E-170 |
| Measured | Predicted | 0.0816 (0.0056) | 4.27E-48 | Measured<br>Observed | 0.858<br>(0.0222) | <1.0E-200 |
| **AUDIT-Problems** |  |  |  |  |  |  |
| Measured | Observed | 0.0468 (0.0049) | 1.28E-21 | Unmeasured<br>Predicted | 0.9126<br>(0.0583) | 2.90E-55 |
| Unmeasured | Predicted | 0.0647 (0.0034) | 9.73E-81 | Measured<br>Predicted | 0.9627<br>(0.0381) | 5.30E-141 |
| Measured | Predicted | 0.0769 (0.0056) | 6.52E-43 | Measured<br>Observed | 0.7915<br>(0.0387) | 5.0E-93 |

Using LDSC, we estimated heritabilities for men and women combined across the three outcome sets. As shown in **Table 1(a) and Figure 7(a)**, these heritabilities ranged from 0.0468 - 0.0869 in the measured sets to 0.0769 - 0.0843 in the predicted sets and the estimates in the measured sets are similar to those previously reported for measured AUDIT in the UKB^27^. The LDSC estimates are slightly lower than the GCTA estimates which is expected since GCTA estimates the genetic relationship matrix (GRM) directly from individual level genotypes while LDSC estimates the GRM from GWAS summary statistics and an LD reference panel resulting in less precision. Heritability in observed and predicted AUDIT-Total and AUDIT-C while moderate (0.0811 - 0.0843 and 0.0869 - 0.0816, respectively) are similar and highly significant, while the point estimate for observed AUDIT-P (0.0468) is lower than that in predicted AUDIT-P (0.0769).

**Figure 7:**
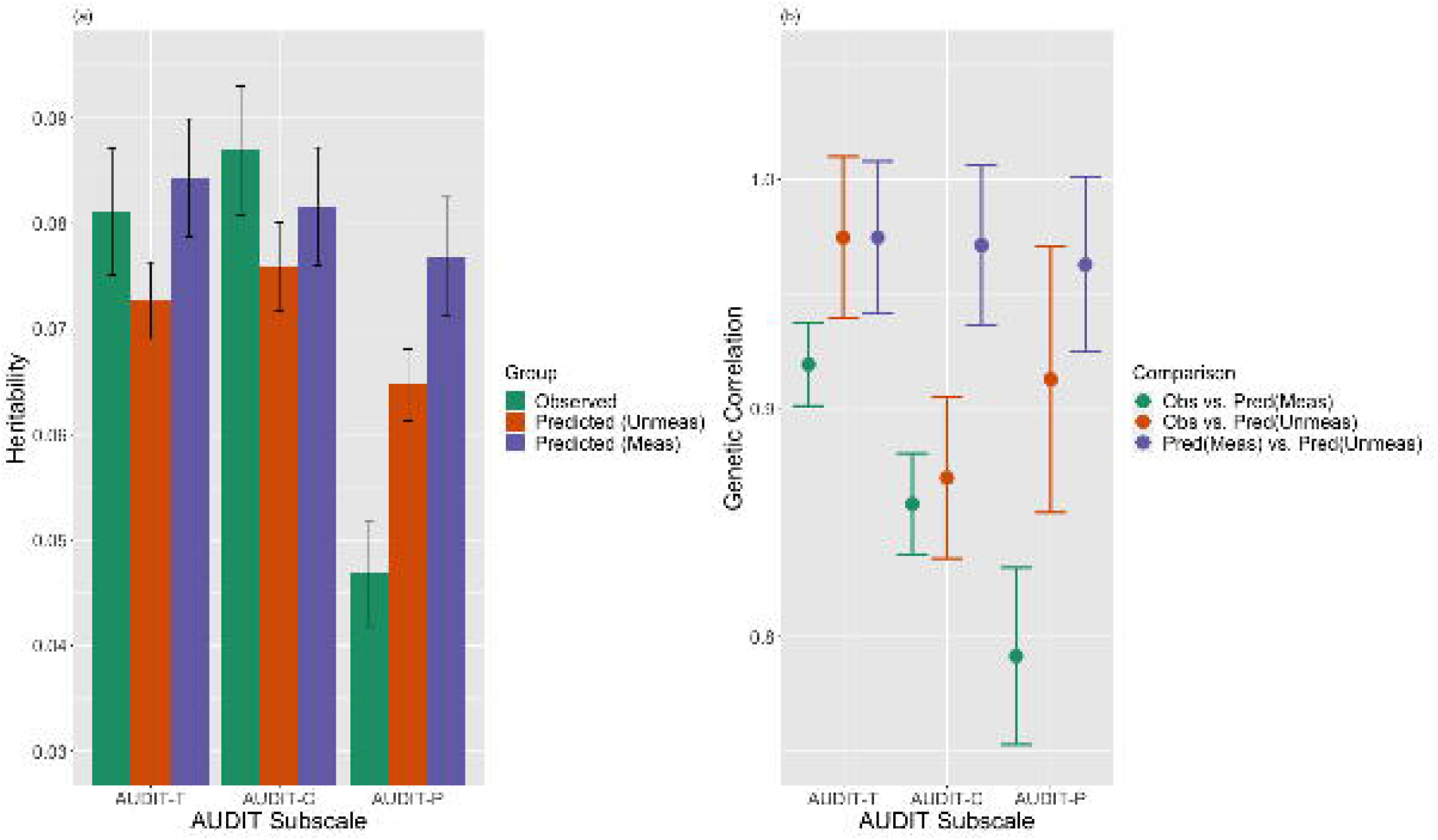
**(a)** LDSC estimated heritabilities. SNP-based heritability estimates for the observed (green) and predicted in the measured (purple) and unmeasured (orange) sets for the AUDIT outcomes. **(b)** LDSC estimated genetic correlations. Genetic correlation estimated between the observed data and predicted scores in the measured sets (green,) the observed data and the predicted scores in the unmeasured sets (orange,) and the predicted scores in the measured and unmeasured sets (purple.)

### Genetic Correlations

Using GCTA and only subjects with measured AUDIT, the *r*_*g*_ between the observed and predicted AUDIT-T was 0.863 (se 0.040) in men and 0.884 (se 0.032) in women. The LDSC-estimated *r*_*g*_ (**Table 1(b) and Figure 7(b))** between observed and predicted AUDIT in the measured set provides an indicator of prediction performance, with *r*_*g*_ between these sets of 0.919 (AUDIT-Total), 0.858 (AUDIT-C), and 0.792 (AUDIT-P.)

## Discussion

The results of our simulation study demonstrated how even moderate levels of random and block-wise missingness rapidly diminish the number of complete cases available in a dataset and the necessity of an innovative approach for prediction analyses in the presence of such missingness. The goal of this methodological work was to extend an ML procedure which could predict missing variables (1) accurately, (2) in an interpretable manner, and (3) in a generalizable framework, (4) using existing software, and (5) for application in biobank-scale datasets with block-wise missing data structures. Our MAGIC-LASSO approach achieves these goals, as demonstrated through the prediction of the AUDIT measures in the UK Biobank study. The consistently high phenotypic and genetic correlations across the observed and predicted sets indicates that the ML procedure is capable of predicting the missing variable with high accuracy and in a manner which faithfully reflects the underlying genetic contribution to the phenotype. Estimated heritabilities in the predicted AUDIT sets were consistent with those in the measured sets and with those previously published. It is further noteworthy to mention that in ML practice, predictive performance in the full set often overestimates the real-life potential of the algorithm to predict missing values. However, our predictive performance in the full set was nearly identical to that in the hold-out test set in the UKB AUDIT application, with a difference in phenotypic correlation of no more than 0.02 between the full and hold-out sets in all three AUDIT measures.

Prediction was less accurate, as measured both by phenotypic correlation and by *r*_*g*_, in the AUDIT-P outcome as compared to AUDIT-Total and AUDIT-C. This demonstrates two considerations, first, that the distribution of the outcome can affect its prediction. Where observations are highly skewed and less evenly distributed across the potential range, prediction is rendered more difficult. Second, prediction performance varies based on the phenotype and the dataset, as observed with the AUDIT measures. The available phenotypes in UKB, in aggregate, lend better information to the prediction of AUDIT-Total and AUDIT-C than of AUDIT-P, although expansion of the variables entering the MAGIC-LASSO model may improve the prediction of AUDIT-P. Furthermore, **Supplementary Figure 3** demonstrates the differing architecture of the predicted scores in Total and Consumption versus Problems, where the first few variables comprise the bulk of the prediction for AUDIT-T and AUDIT-C, while the prediction of AUDIT-P is composed of more variables of small effect. From an epidemiological perspective, it is also noteworthy to consider that problematic alcohol use, as measured across multiple behaviors, is a more complex, and therefore more difficult to predict phenotype than quantity of alcohol consumed, which may explain why the genetic correlation between the observed and predicted AUDIT-P was lower than seen in AUDIT-Total and AUDIT-C.

Strengths of the MAGIC-LASSO include, first, it can be applied using existing packages in the R software environment. Second, the prediction process is straightforward and transparent. The MAGIC-LASSO is built on the foundation of the Group-LASSO, a statistically rigorous framework with well-established properties which allow the user access to the regression structure of the prediction. Third, it is applicable to large biobank-scale environments where missing-at-random structures cannot be assumed. The application of the MAGIC-LASSO for variable imputation can confer great power gains for genetic analyses, as demonstrated using AUDIT in UKB. AUDIT and genotypic data were directly measured in 117,559 European ancestry individuals in the UKB sample. Predicting AUDIT in the unmeasured subjects added 242,421 independent samples for downstream GWAS, representing a 56% increase in effective sample size. Finally, the MAGIC-LASSO is a flexible framework allowing for straightforward adaptations for application to datasets of various structures and outcomes of different characteristics.

Limitations of the MAGIC-LASSO framework include the limitations of the Group-LASSO procedure to account for interaction effects of covariates. The current demonstrated implementation of the approach is also limited to the linear regression framework. As a novel application, rather than a novel algorithm, the approach is intended to serve as a tool which may be readily modified to varying applications, such as a prediction of binary or multi-category outcomes. In fact, the Group-LASSO function employed (*grpreg*) is applicable to many other types of outcomes. Predicting missing outcomes in the presence of missing predictors, using the final Group-LASSO iteration of the MAGIC-LASSO introduces some bias to the predictors, given that these predictors must be dropped from the regression equation. In the presence of large sample sizes, such as those typically collected in biobanks however, this bias and associated loss of prediction accuracy is somewhat mitigated. Both simulation and the UKB application examples demonstrate that the predicted outcomes were overall robust to this missingness.

Furthermore, the application of the MAGIC-LASSO also requires some degree of statistical judgment to be rendered during model-fitting, including the selection of the cutpoint during the clustering step and the stopping criteria for desired model parsimony. In this way, the approach is decidedly non “black box” in nature and requires the researcher to interact with the modeling process. Future directions in our research aim to implement missingness adaptation approaches into additional ML paradigms, such as boosting and random forests.

Despite these limitations, the method demonstrated strong predictive performance, both in simulations and in the real data application in UKB and represents an innovative contribution to the field of epidemiological research in biobanks. The method is accessible through open-source software and transparent in nature, allowing the user to assess performance and understand the full regression procedure constructing the predicted outcomes. The MAGIC-LASSO is an additional tool now available to researchers to further harness the discovery potential inherent in large data collections and maximize the return on the financial and altruistic participant time and effort contributions invested in the assembly and management of biobank resources.

## Acknowledgements

This research has been conducted using the UK Biobank Resource application number 30782. This work was supported by the National Institutes of Health [grant number P50AA022537; grant number T32MH020030, to [AEG]; and grant number K01MH113848, to [REP]] and The Brain & Behavior Research Foundation [grant number 28632 NARSAD P&S Fund, to [REP]].

## Author Contributions

All authors contributed to the concept and design of the study. REP. and BTW. contributed to the data acquisition. All authors contributed to the analyses, interpretation of the data and the results, as well as to substantive revisions of the manuscript.

## Data Availability

The UKB data utilized in this research is available to, “bona fide researchers for health-related research in the public interest^28^,” through an application process accessible through the UKB website, https://www.ukbiobank.ac.uk/.

## Additional Information

The authors have no competing interests to disclose.

