## Supplemental Material for "Missingness Adapted Group Informed Clustered (MAGIC)-LASSO: A novel paradigm for prediction in data with widespread non-random missingness"

**Contents:**

Supplementary Section 1

LASSO background

Group-LASSO background

Supplementary Section 2

Simulation study details

Supplementary Table 1

Missingness simulation results

Supplementary Tables 2-7

MAGIC-LASSO simulation results

Supplementary Table 8

Clustering details

Supplementary Table 9

Step-wise Group-LASSO results

Supplementary Table 10

GCTA heritabilities and genetic correlation estimates

Supplementary Figure 1

UKB filtering

Supplementary Figure 2

Step-wise model results

**Supplementary Section 1**

**LASSO background:**

As a member of the family of penalized regression ML techniques, the LASSO^1^ is well established and popular. Often presented in the context of the elastic net^2^ formulation, the general formula for the LASSO may be found under the linear regression paradigm by estimating the values which minimize:

$\frac{1}{2N}\sum_{i=1}^{N} \left( y_{i}-\beta_{0}-x_{i}^{T}\beta\right)^{2}+\lambda\left[ \frac{\left( 1-\alpha\right){\|\beta\|}_{2}^{2}}{2}+\alpha{\|\beta\|}_{1} \right]$,

where there are $i=1,\ldots,N$ total participants, $y_{i}$ represents outcome observation $i$ and $x_{i}^{T}$ is a vector of predictors. The $\lambda$ is a tuning parameter that adjusts the amount of shrinkage (penalization) applied to the model and $\alpha$ mixes the $L_{1}$norm and $L_{2}$norm penalties, where:

$L_{1} penalty, {\|\beta\|}_{1}=\sum_{i=1}^{N} \left| x_{i} \right|$, and

$L_{2} penalty, {\|\beta\|}_{2}^{2}=\sum_{i=1}^{N} \left| x_{i} \right|^{2}$,

such that when $\alpha=1$, the formula becomes the LASSO and when $\alpha=0$, the formula becomes ridge regression^3^; mixing the two parameters using $0<\alpha<1$ results in the classic elastic net design. The nature of the ridge penalty prevents any coefficient estimate to shrink to exactly zero, making it more useful for addressing multicollinearity than dimension reduction, while the LASSO encourages sparsity and the coefficients of some covariates are allowed to shrink to exactly zero, making it a useful tool for variable selection. The mixing parameter is generally chosen by the user and set for the duration of the experiment. Where the overarching goal is to identify a parsimonious set of covariates from a large pool which may accurately approximate an outcome, setting $\alpha=1$ is generally appropriate because the LASSO is well suited to achieve this. The tuning parameter ($\lambda$) is best chosen through cross-validation, a method by which a single observation is held out during the fitting process and the resulting model used to predict the outcome of the left-out observation. Repeating this process for every observation in the dataset allows for the calculation of a mean error rate of prediction. The lambda associated with the model with the smallest mean error rate of prediction is generally the best model^4^.

**Group LASSO background:**

In traditional LASSO, categorical covariates can be included by coding them according to a numerical scale. This is not ideal, as it assumes equal spacing between and inherent ordering within categories^5^. A solution to this is to dummy-code the $k$-level categorical measures by augmenting the representation of a single variable into $k-1$ binary variables. However, traditional LASSO treats each of these as individual measures which may result in shrinkage of some but not all categories within a single covariate, rendering interpretation of selected variables difficult. The Group LASSO^6^ was developed to encourage sparsity at the factor level, such that all categories of a given variable are included or excluded from the model as a set. For samples j=1…J, the Group LASSO is found as the solution to:

$\frac{1}{2}{\|Y-\sum_{j=1}^{J} X_{i}\beta_{j}\|}^{2}+\lambda\sum_{j=1}^{J} {\|\beta_{j}\|}_{K_{j}}$,

Where ${\|\beta_{j}\|}_{K_{j}}=\left( \beta_{j}^{T}K_{j}\beta_{j} \right)^{\frac{1}{2}}$.

In the simplest form, $K_{j}$ can be an identity matrix, but it may take a variety of forms. To fit a model within this framework, each categorical variable must be expanded into binary dummy-coded columns.

**Supplementary Section 1 References:**

1.	Tibshirani, R. Regression Shrinkage and Selection via the Lasso. *J. R. Stat. Soc. Series B Stat. Methodol.* **58**, 267–288 (1996).

2.	Zou, H. & Hastie, T. Regularization and variable selection via the elastic net. *J. R. Stat. Soc. Series B Stat. Methodol.* **67**, 301–320 (04/2005).

3.	Hoerl, A. E. & Kennard, R. W. Ridge Regression: Biased Estimation for Nonorthogonal Problems. *Technometrics* **12**, 55–67 (02/1970).

4.	Hastie, T., Tibshirani, R. & Friedman, J. *The elements of statistical learning*. (Springer, 2009).

5.	*Applied linear statistical models*. (McGraw-Hill Irwin, 2005).

6.	Yuan, M. & Lin, Y. Model selection and estimation in regression with grouped variables. *J. R. Stat. Soc. Series B Stat. Methodol.* **68**, 49–67 (02/2006).

**Supplementary Section 2**

**Simulation study details:**

We conducted a series of simulations, under 6 separate sets of conditions and parameters, over 100 iterations each. For each set of conditions, the following models were run:

1. LASSO (treating categorical covariates as numeric) and Group-LASSO with no missing data
2. LASSO and Group-LASSO with only response missingness, ie, no missingness in the predictor space
3. LASSO and Group-LASSO with response missingness and 5, 10, and 15% random missingness in the predictor space
4. LASSO, Group-LASSO, and MAGIC-LASSO with response missingness, 5% random missingness, and blockwise missingness, with some complete cases present in the predictor space
5. MAGIC-LASSO with response missingness, 5% random missingness, and blockwise missingness, with no complete cases present in the predictor space, using no hold-out set, a 5% hold-out set before the first Group-LASSO iteration, or a 5% hold-out set before the final Group-LASSO iteration only

We utilized 6 separate condition sets, with the following parameters:

| **Set** | **N** | **P** | **Cat Lev** | **Beta Inc** | **Prop Comp** | **Int** | **N Beta** | **Betas** | **Cuts** |
| --- | --- | --- | --- | --- | --- | --- | --- | --- | --- |
| 1 | 10000 | 200 | 4 | (1, -4, 3) | 0.1 | -10 | 10 | (0.5,2,-2,5,5,-5,7,7,7,-7) | (6,7,8,9,10,11,12) |
| 2 | 50000 | 250 | 6 | (1, -4, 3, 0.5, -2) | 0.005 | -10 | 10 | (0.5,2,-2,5,5,-5,7,7,7,-7) | (3, 4, 5, 6,7,8,9,10,11,12) |
| 3 | 50000 | 250 | 6 | (1, -4, 3, 0.5, -2) | 0.005 | -10 | 10 | (0.5,0.5,-0.5,-0.5,0.75,  -0.75,1,1,-1,-1) | (3, 4, 5, 6,7,8,9,10,11,12) |
| 4 | 50000 | 250 | 6 | (1, -4, 3, 0.5, -2) | 0.005 | -10 | 5 | (0.5, -0.5,0.75,1,-1) | (3, 4, 5, 6,7,8,9,10,11,12) |
| 5 | 50000 | 250 | 6 | (1, -4, 3, 0.5, -2) | 0.005 | -10 | 20 | (0.5,0.5,0.5,-0.5,-0.5,0.75,  -0.75,0.05,0.1,0.15,1,1,-1,  -1,-1,2,-2,3,3,-3) | (3, 4, 5, 6,7,8,9,10,11,12) |
| 6 | 50000 | 250 | 6 | (1.25, -2, 2, -1.25, 0.25) | 0.005 | -10 | 10 | (0.5,0.5,-0.5,-0.5,0.75,  -0.75,1,1,-1,-1) | (3, 4, 5, 6,7,8,9,10,11,12) |

Legend:

Set: Simulation set

N: Sample size

P: Number of predictors

Cat Lev: Number of categories for categorical predictors

Beta Inc: Increments of increase/decrease for each level of categorical variables from the reference level; used for assigning beta values to dummy-coded categorical variables

Prop Comp: Proportion of the data with complete cases

Int: Intercept

N Beta: Number of true betas

Betas: True beta values

Cuts: Cutpoints for the hierarchical clustering trees

**Supplementary Table 1**

| **Supplementary Table 1:** Simulation results showing number of complete cases as a function of a range random missingness and block-wise missingness across 100 iterations. | | | | | | | | | | | | | |
| --- | --- | --- | --- | --- | --- | --- | --- | --- | --- | --- | --- | --- | --- |
| **Percent**  **Rand.** | **No.**  **Blocks** | **Mean**  **CC** | **Median**  **CC** | **Min.**  **CC** | **Max.**  **CC** | **Mean**  **Total**  **Miss** | **Median**  **Total**  **Miss** | **Min.**  **Total**  **Miss** | **Max.**  **Total**  **Miss** | **Mean**  **Miss**  **Percent** | **Median**  **Miss**  **Percent** | **Min.**  **Miss**  **Percent** | **Max.**  **Miss**  **Percent** |
| 1 | 0 | 1352 | 1353 | 1274 | 1436 | 19900 | 19900 | 19860 | 19925 | 0.01 | 0.01 | 0.01 | 0.01 |
| 1 | 5 | 1309 | 1308 | 1229 | 1384 | 23058 | 22840 | 20592 | 27305 | 0.012 | 0.011 | 0.01 | 0.014 |
| 1 | 10 | 1310 | 1314 | 1220 | 1395 | 22972 | 22486 | 20585 | 27329 | 0.011 | 0.011 | 0.01 | 0.014 |
| 1 | 15 | 1310 | 1310 | 1212 | 1401 | 22995 | 22868 | 20483 | 26832 | 0.011 | 0.011 | 0.01 | 0.013 |
| 1 | 20 | 1311 | 1310 | 1223 | 1382 | 22931 | 22866 | 20397 | 26604 | 0.011 | 0.011 | 0.01 | 0.013 |
| 2 | 0 | 183 | 184 | 148 | 218 | 39604 | 39606 | 39550 | 39654 | 0.02 | 0.02 | 0.02 | 0.02 |
| 2 | 5 | 177 | 177 | 141 | 211 | 42722 | 42519 | 40262 | 46950 | 0.021 | 0.021 | 0.02 | 0.023 |
| 2 | 10 | 177 | 177 | 145 | 209 | 42637 | 42169 | 40288 | 46916 | 0.021 | 0.021 | 0.02 | 0.023 |
| 2 | 15 | 177 | 177 | 144 | 212 | 42651 | 42556 | 40198 | 46453 | 0.021 | 0.021 | 0.02 | 0.023 |
| 2 | 20 | 178 | 177 | 141 | 215 | 42593 | 42530 | 40065 | 46270 | 0.021 | 0.021 | 0.02 | 0.023 |
| 3 | 0 | 25 | 26 | 15 | 38 | 59112 | 59115 | 59030 | 59185 | 0.03 | 0.03 | 0.03 | 0.03 |
| 3 | 5 | 25 | 25 | 15 | 36 | 62159 | 61936 | 59778 | 66390 | 0.031 | 0.031 | 0.03 | 0.033 |
| 3 | 10 | 24 | 25 | 15 | 38 | 62078 | 61633 | 59724 | 66418 | 0.031 | 0.031 | 0.03 | 0.033 |
| 3 | 15 | 25 | 25 | 15 | 36 | 62152 | 62040 | 59658 | 65921 | 0.031 | 0.031 | 0.03 | 0.033 |
| 3 | 20 | 25 | 25 | 15 | 35 | 62027 | 61966 | 59556 | 65678 | 0.031 | 0.031 | 0.03 | 0.033 |
| 4 | 0 | 4 | 3 | 0 | 8 | 78427 | 78431 | 78303 | 78518 | 0.039 | 0.039 | 0.039 | 0.039 |
| 4 | 5 | 3 | 3 | 0 | 8 | 81394 | 81131 | 79156 | 85674 | 0.041 | 0.041 | 0.04 | 0.043 |
| 4 | 10 | 3 | 3 | 0 | 8 | 81371 | 80922 | 79080 | 85589 | 0.041 | 0.04 | 0.04 | 0.043 |
| 4 | 15 | 3 | 3 | 0 | 8 | 81429 | 81312 | 79026 | 85156 | 0.041 | 0.041 | 0.04 | 0.043 |
| 4 | 20 | 3 | 3 | 0 | 8 | 81325 | 81237 | 78882 | 84930 | 0.041 | 0.041 | 0.039 | 0.042 |
| 5 | 0 | 1 | 0 | 0 | 3 | 97547 | 97552 | 97417 | 97665 | 0.049 | 0.049 | 0.049 | 0.049 |
| 5 | 5 | 1 | 0 | 0 | 3 | 100513 | 100198 | 98258 | 104719 | 0.05 | 0.05 | 0.049 | 0.052 |
| 5 | 10 | 1 | 0 | 0 | 3 | 100421 | 99966 | 98171 | 104737 | 0.05 | 0.05 | 0.049 | 0.052 |
| 5 | 15 | 1 | 0 | 0 | 3 | 100517 | 100439 | 98124 | 104193 | 0.05 | 0.05 | 0.049 | 0.052 |
| 5 | 20 | 1 | 0 | 0 | 3 | 100405 | 100377 | 97953 | 104060 | 0.05 | 0.05 | 0.049 | 0.052 |
| 6 | 0 | 0 | 0 | 0 | 1 | 116477 | 116478 | 116321 | 116651 | 0.058 | 0.058 | 0.058 | 0.058 |
| 6 | 5 | 0 | 0 | 0 | 1 | 119448 | 119230 | 117137 | 123429 | 0.06 | 0.06 | 0.059 | 0.062 |
| 6 | 10 | 0 | 0 | 0 | 1 | 119307 | 118946 | 117126 | 123515 | 0.06 | 0.059 | 0.059 | 0.062 |
| 6 | 15 | 0 | 0 | 0 | 1 | 119417 | 119312 | 116983 | 123147 | 0.06 | 0.06 | 0.058 | 0.062 |
| 6 | 20 | 0 | 0 | 0 | 1 | 119274 | 119155 | 116931 | 122792 | 0.06 | 0.06 | 0.058 | 0.061 |
| 7 | 0 | 0 | 0 | 0 | 1 | 135218 | 135218 | 135047 | 135414 | 0.068 | 0.068 | 0.068 | 0.068 |
| 7 | 5 | 0 | 0 | 0 | 1 | 138145 | 137890 | 135765 | 142296 | 0.069 | 0.069 | 0.068 | 0.071 |
| 7 | 10 | 0 | 0 | 0 | 1 | 138021 | 137626 | 135973 | 142202 | 0.069 | 0.069 | 0.068 | 0.071 |
| 7 | 15 | 0 | 0 | 0 | 1 | 138105 | 138032 | 135814 | 141844 | 0.069 | 0.069 | 0.068 | 0.071 |
| 7 | 20 | 0 | 0 | 0 | 1 | 137972 | 137920 | 135693 | 141438 | 0.069 | 0.069 | 0.068 | 0.071 |
| 8 | 0 | 0 | 0 | 0 | 1 | 153776 | 153776 | 153574 | 153966 | 0.077 | 0.077 | 0.077 | 0.077 |
| 8 | 5 | 0 | 0 | 0 | 1 | 156680 | 156462 | 154325 | 160616 | 0.078 | 0.078 | 0.077 | 0.08 |
| 8 | 10 | 0 | 0 | 0 | 1 | 156564 | 156159 | 154464 | 160792 | 0.078 | 0.078 | 0.077 | 0.08 |
| 8 | 15 | 0 | 0 | 0 | 1 | 156654 | 156516 | 154325 | 160347 | 0.078 | 0.078 | 0.077 | 0.08 |
| 8 | 20 | 0 | 0 | 0 | 1 | 156493 | 156518 | 154122 | 159922 | 0.078 | 0.078 | 0.077 | 0.08 |
| 9 | 0 | 0 | 0 | 0 | 1 | 172150 | 172151 | 171923 | 172362 | 0.086 | 0.086 | 0.086 | 0.086 |
| 9 | 5 | 0 | 0 | 0 | 1 | 175041 | 174864 | 172827 | 179026 | 0.088 | 0.087 | 0.086 | 0.09 |
| 9 | 10 | 0 | 0 | 0 | 1 | 174911 | 174531 | 172812 | 179024 | 0.087 | 0.087 | 0.086 | 0.09 |
| 9 | 15 | 0 | 0 | 0 | 1 | 175050 | 174900 | 172558 | 178780 | 0.088 | 0.087 | 0.086 | 0.089 |
| 9 | 20 | 0 | 0 | 0 | 1 | 174798 | 174707 | 172376 | 178361 | 0.087 | 0.087 | 0.086 | 0.089 |
| 10 | 0 | 0 | 0 | 0 | 1 | 190340 | 190344 | 190081 | 190645 | 0.095 | 0.095 | 0.095 | 0.095 |
| 10 | 5 | 0 | 0 | 0 | 1 | 193247 | 193007 | 190797 | 197206 | 0.097 | 0.097 | 0.095 | 0.099 |
| 10 | 10 | 0 | 0 | 0 | 1 | 193063 | 192717 | 190750 | 197200 | 0.097 | 0.096 | 0.095 | 0.099 |
| 10 | 15 | 0 | 0 | 0 | 1 | 193184 | 193069 | 190930 | 196650 | 0.097 | 0.097 | 0.095 | 0.098 |
| 10 | 20 | 0 | 0 | 0 | 1 | 192920 | 192930 | 190625 | 196482 | 0.096 | 0.096 | 0.095 | 0.098 |
| 11 | 0 | 0 | 0 | 0 | 0 | 208344 | 208352 | 208083 | 208642 | 0.104 | 0.104 | 0.104 | 0.104 |
| 11 | 5 | 0 | 0 | 0 | 0 | 211223 | 211045 | 208897 | 215043 | 0.106 | 0.106 | 0.104 | 0.108 |
| 11 | 10 | 0 | 0 | 0 | 0 | 211034 | 210664 | 208779 | 215406 | 0.106 | 0.105 | 0.104 | 0.108 |
| 11 | 15 | 0 | 0 | 0 | 0 | 211162 | 211027 | 208824 | 214866 | 0.106 | 0.106 | 0.104 | 0.107 |
| 11 | 20 | 0 | 0 | 0 | 0 | 210883 | 210768 | 208544 | 214444 | 0.105 | 0.105 | 0.104 | 0.107 |

Legend:

Percent Rand.: Percent random missingness

No. Blocks: Number of missingness blocks

Mean CC: Mean number of complete cases

Median CC: Median number of complete cases

Min. CC: Minimum number of complete cases

Max. CC: Maximum number of complete cases

Mean Total Miss: Mean total number of missing observations

Median Total Miss: Median total number of missing observations

Min. Total Miss: Minimum number of missing observations

Max. Total Miss: Maximum number of missing observations

Mean Miss Percent: Mean percent missing observations

Median Miss Percent: Median percent missing observations

Min. Miss Percent: Minimum percent missing observations

Max. Miss Percent: Maximum percent missing observations

**Supplementary Table 2:** Simulation results for MAGIC-LASSO simulation, scenario 1.

| **Scenario 1** | COR | COR (All) | MSE | MSE (All) | N-TRAIN | Miss Prop |
| --- | --- | --- | --- | --- | --- | --- |
| Group-LASSO  No missingness | 0.9948 | 0.9948 | 4.01 | 3.97 | 10000 | 0 |
| Group-LASSO | 0.7614 | 0.9646 | 495.93 | 31.69 | 870 | 0.123 |
| MAGIC-LASSO (n trees = 6) | 0.7602 | 0.965 | 506.72 | 31.87 | 944 | 0.123 |
| MAGIC-LASSO (n trees = 7) | 0.7602 | 0.9648 | 504.38 | 31.83 | 944 | 0.123 |
| MAGIC-LASSO (n trees = 8) | 0.7602 | 0.9649 | 505.05 | 31.81 | 944 | 0.123 |
| MAGIC-LASSO (n trees = 9) | 0.7603 | 0.9649 | 503.94 | 31.77 | 946 | 0.123 |
| MAGIC-LASSO (n trees = 10) | 0.7601 | 0.9649 | 504.51 | 31.78 | 946 | 0.123 |
| MAGIC-LASSO (n trees = 11) | 0.7597 | 0.9649 | 502.39 | 31.68 | 924 | 0.123 |
| MAGIC-LASSO (n trees = 12) | 0.7595 | 0.9648 | 504.13 | 31.74 | 924 | 0.123 |
| MAGIC-LASSO (n trees = 6) | 0.5937 | 0.65 | 555.6 | 460.57 | 3401 | 0.123 |
| MAGIC-LASSO (n trees = 7) | 0.6217 | 0.6768 | 519.32 | 426.33 | 3129 | 0.136 |
| MAGIC-LASSO (n trees = 8) | 0.6308 | 0.6859 | 502.3 | 414.35 | 3014 | 0.136 |
| MAGIC-LASSO (n trees = 9) | 0.6112 | 0.6616 | 554.1 | 464.06 | 3397 | 0.136 |
| MAGIC-LASSO (n trees = 10) | 0.5646 | 0.622 | 616.04 | 514.65 | 3865 | 0.136 |
| MAGIC-LASSO (n trees = 11) | 0.4889 | 0.5493 | 717.01 | 634.64 | 3919 | 0.136 |
| MAGIC-LASSO (n trees = 12) | 0.5094 | 0.5696 | 686.76 | 604.67 | 3759 | 0.136 |

Legend:

COR: correlation between observed and predicted outcomes for the missing outcomes

COR-All: correlation between observed and predicted outcomes for the full data set

MSE: mean squared error of prediction between observed and predicted outcomes for the missing outcomes

MSE-All: mean squared error of prediction between observed and predicted outcomes for the full dataset

N-Train: number of observations used to construct the model

Miss Prop: proportion of the total data set missing

n trees: number of tree utilized for cutting the clustering trees

*Shaded cells indicate scenarios with zero complete data cases

**Supplementary Table 3:** Simulation results for MAGIC-LASSO simulation, scenario 2.

| **Scenario 2** | COR | COR-All | MSE | MSE-all | N-TRAIN | Miss Prop |
| --- | --- | --- | --- | --- | --- | --- |
| Group-LASSO  No missingness | 0.994 | 0.9942 | 3.96 | 3.98 | 50000 | 0 |
| Group-LASSO | 0.7037 | 0.9035 | 565.3 | 116.94 | 218 | 0.140 |
| MAGIC-LASSO (n trees = 3) | 0.7737 | 0.9565 | 372.89 | 54.99 | 747 | 0.140 |
| MAGIC-LASSO (n trees = 4) | 0.7817 | 0.9554 | 386.52 | 58.32 | 541 | 0.140 |
| MAGIC-LASSO (n trees = 5) | 0.7975 | 0.9483 | 382.6 | 72.7 | 350 | 0.140 |
| MAGIC-LASSO (n trees = 6) | 0.7967 | 0.9491 | 381.23 | 72.4 | 327 | 0.140 |
| MAGIC-LASSO (n trees = 7) | 0.7849 | 0.9498 | 387.7 | 71.42 | 391 | 0.140 |
| MAGIC-LASSO (n trees = 8) | 0.7818 | 0.9468 | 388.6 | 71.77 | 389 | 0.140 |
| MAGIC-LASSO (n trees = 9) | 0.7844 | 0.9493 | 384.03 | 68.59 | 465 | 0.140 |
| MAGIC-LASSO (n trees = 10) | 0.7915 | 0.9481 | 371.38 | 71.16 | 354 | 0.140 |
| MAGIC-LASSO (n trees = 11) | 0.7835 | 0.9461 | 372.2 | 74.79 | 335 | 0.140 |
| MAGIC-LASSO (n trees = 12) | 0.7837 | 0.9462 | 371.99 | 74.69 | 332 | 0.140 |
| MAGIC-LASSO (n trees = 3) | 0.7227 | 0.7602 | 366.25 | 300.17 | 8778 | 0.141 |
| MAGIC-LASSO (n trees = 4) | 0.6926 | 0.7306 | 398.93 | 333.91 | 9153 | 0.141 |
| MAGIC-LASSO (n trees = 5) | 0.666 | 0.7088 | 442.87 | 363.99 | 10258 | 0.141 |
| MAGIC-LASSO (n trees = 6) | 0.6463 | 0.6898 | 475.45 | 388.76 | 10720 | 0.141 |
| MAGIC-LASSO (n trees = 7) | 0.6468 | 0.6903 | 485.82 | 399.66 | 10509 | 0.141 |
| MAGIC-LASSO (n trees = 8) | 0.6681 | 0.7079 | 431.07 | 371.34 | 10099 | 0.141 |
| MAGIC-LASSO (n trees = 9) | 0.6634 | 0.7001 | 461.06 | 410.52 | 10058 | 0.141 |
| MAGIC-LASSO (n trees = 10) | 0.6107 | 0.6524 | 523.93 | 471.25 | 12358 | 0.141 |
| MAGIC-LASSO (n trees = 11) | 0.6045 | 0.6468 | 532.3 | 478.18 | 12562 | 0.141 |
| MAGIC-LASSO (n trees = 12) | 0.6171 | 0.6575 | 503.32 | 450.27 | 12047 | 0.141 |

Legend:

COR: correlation between observed and predicted outcomes for the missing outcomes

COR-All: correlation between observed and predicted outcomes for the full data set

MSE: mean squared error of prediction between observed and predicted outcomes for the missing outcomes

MSE-All: mean squared error of prediction between observed and predicted outcomes for the full dataset

N-Train: number of observations used to construct the model

Miss Prop: proportion of the total data set missing

n trees: number of tree utilized for cutting the clustering trees

*Shaded cells indicate scenarios with zero complete data cases

**Supplementary Table 4:** Simulation results for MAGIC-LASSO simulation, scenario 3.

| **Scenario 3** | COR | COR-All | MSE | MSE-all | N-TRAIN | Miss Prop |
| --- | --- | --- | --- | --- | --- | --- |
| Group-LASSO  No missingness | 0.9682 | 0.968 | 3.97 | 3.99 | 50000 | 0 |
| Group-LASSO | 0.6537 | 0.7985 | 95.41 | 39.11 | 218 | 0.140 |
| MAGIC-LASSO (n trees = 3) | 0.7554 | 0.9055 | 36.73 | 14.5 | 1050 | 0.140 |
| MAGIC-LASSO (n trees = 4) | 0.7516 | 0.902 | 45.85 | 15.66 | 825 | 0.140 |
| MAGIC-LASSO (n trees = 5) | 0.7521 | 0.895 | 46.08 | 19.28 | 439 | 0.140 |
| MAGIC-LASSO (n trees = 6) | 0.7466 | 0.89 | 47.41 | 19.34 | 392 | 0.140 |
| MAGIC-LASSO (n trees = 7) | 0.7462 | 0.8913 | 49.43 | 21.8 | 477 | 0.140 |
| MAGIC-LASSO (n trees = 8) | 0.738 | 0.8839 | 49.04 | 22.37 | 483 | 0.140 |
| MAGIC-LASSO (n trees = 9) | 0.7382 | 0.8842 | 50.89 | 24.03 | 580 | 0.140 |
| MAGIC-LASSO (n trees = 10) | 0.737 | 0.888 | 46.5 | 17.86 | 482 | 0.140 |
| MAGIC-LASSO (n trees = 11) | 0.7376 | 0.8815 | 48.9 | 19.91 | 445 | 0.140 |
| MAGIC-LASSO (n trees = 12) | 0.737 | 0.8808 | 49.59 | 20.03 | 437 | 0.140 |
| MAGIC-LASSO (n trees = 3) | 0.683 | 0.7161 | 31.28 | 28.15 | 9506 | 0.141 |
| MAGIC-LASSO (n trees = 4) | 0.6649 | 0.6962 | 44.02 | 39.66 | 8628 | 0.141 |
| MAGIC-LASSO (n trees = 5) | 0.6456 | 0.6811 | 39.85 | 36.33 | 9241 | 0.141 |
| MAGIC-LASSO (n trees = 6) | 0.6172 | 0.653 | 64.98 | 61.59 | 9340 | 0.141 |
| MAGIC-LASSO (n trees = 7) | 0.6251 | 0.6614 | 50.13 | 43.4 | 9478 | 0.141 |
| MAGIC-LASSO (n trees = 8) | 0.6207 | 0.6564 | 56.37 | 45.27 | 9513 | 0.141 |
| MAGIC-LASSO (n trees = 9) | 0.6107 | 0.6422 | 42.27 | 38.64 | 9392 | 0.141 |
| MAGIC-LASSO (n trees = 10) | 0.5882 | 0.6152 | 45.55 | 43.59 | 10250 | 0.141 |
| MAGIC-LASSO (n trees = 11) | 0.5814 | 0.6118 | 47.13 | 43.94 | 10647 | 0.141 |
| MAGIC-LASSO (n trees = 12) | 0.5829 | 0.6123 | 51.02 | 47.55 | 10541 | 0.141 |

Legend:

COR: correlation between observed and predicted outcomes for the missing outcomes

COR-All: correlation between observed and predicted outcomes for the full data set

MSE: mean squared error of prediction between observed and predicted outcomes for the missing outcomes

MSE-All: mean squared error of prediction between observed and predicted outcomes for the full dataset

N-Train: number of observations used to construct the model

Miss Prop: proportion of the total data set missing

n trees: number of tree utilized for cutting the clustering trees

*Shaded cells indicate scenarios with zero complete data cases

**Supplementary Table 5:** Simulation results for MAGIC-LASSO simulation, scenario 4.

| **Scenario 4** | COR | COR-All | MSE | MSE-all | N-TRAIN | Miss Prop |
| --- | --- | --- | --- | --- | --- | --- |
| Group-LASSO  No missingness | 0.9541 | 0.9541 | 3.98 | 3.99 | 50000 | 0 |
| Group-LASSO | 0.6632 | 0.823 | 68.64 | 26.35 | 218 | 0.140 |
| MAGIC-LASSO (n trees = 3) | 0.7144 | 0.8759 | 30.53 | 14.22 | 799 | 0.140 |
| MAGIC-LASSO (n trees = 4) | 0.7242 | 0.8816 | 48.19 | 14.12 | 679 | 0.140 |
| MAGIC-LASSO (n trees = 5) | 0.72 | 0.8724 | 71 | 17.22 | 395 | 0.140 |
| MAGIC-LASSO (n trees = 6) | 0.7074 | 0.8615 | 47.54 | 16.83 | 357 | 0.140 |
| MAGIC-LASSO (n trees = 7) | 0.71 | 0.861 | 52.22 | 17.7 | 440 | 0.140 |
| MAGIC-LASSO (n trees = 8) | 0.7073 | 0.8557 | 58.41 | 18.99 | 445 | 0.140 |
| MAGIC-LASSO (n trees = 9) | 0.714 | 0.8633 | 68.88 | 22.32 | 523 | 0.140 |
| MAGIC-LASSO (n trees = 10) | 0.7081 | 0.8663 | 43.07 | 16.24 | 420 | 0.140 |
| MAGIC-LASSO (n trees = 11) | 0.7168 | 0.8729 | 32.86 | 14.6 | 383 | 0.140 |
| MAGIC-LASSO (n trees = 12) | 0.7178 | 0.8738 | 32.89 | 14.51 | 384 | 0.140 |
| MAGIC-LASSO (n trees = 3) | 0.6495 | 0.6818 | 23.41 | 20.9 | 9042 | 0.141 |
| MAGIC-LASSO (n trees = 4) | 0.6296 | 0.6636 | 25.17 | 22.5 | 8351 | 0.141 |
| MAGIC-LASSO (n trees = 5) | 0.5833 | 0.6215 | 39.65 | 35.05 | 8510 | 0.141 |
| MAGIC-LASSO (n trees = 6) | 0.563 | 0.6031 | 37.76 | 33.55 | 9040 | 0.141 |
| MAGIC-LASSO (n trees = 7) | 0.5646 | 0.603 | 39.88 | 35.7 | 8752 | 0.141 |
| MAGIC-LASSO (n trees = 8) | 0.5881 | 0.6244 | 35.36 | 32.15 | 8472 | 0.141 |
| MAGIC-LASSO (n trees = 9) | 0.6215 | 0.6538 | 26.45 | 24.27 | 8403 | 0.141 |
| MAGIC-LASSO (n trees = 10) | 0.6097 | 0.6417 | 26.25 | 24.11 | 9086 | 0.141 |
| MAGIC-LASSO (n trees = 11) | 0.6108 | 0.6432 | 25.87 | 23.7 | 9215 | 0.141 |
| MAGIC-LASSO (n trees = 12) | 0.6105 | 0.6412 | 27.23 | 25.4 | 8917 | 0.141 |

Legend:

COR: correlation between observed and predicted outcomes for the missing outcomes

COR-All: correlation between observed and predicted outcomes for the full data set

MSE: mean squared error of prediction between observed and predicted outcomes for the missing outcomes

MSE-All: mean squared error of prediction between observed and predicted outcomes for the full dataset

N-Train: number of observations used to construct the model

Miss Prop: proportion of the total data set missing

n trees: number of tree utilized for cutting the clustering trees

*Shaded cells indicate scenarios with zero complete data cases

**Supplementary Table 6:** Simulation results for MAGIC-LASSO simulation, scenario 5.

| **Scenario 5** | COR | COR-All | MSE | MSE-all | N-TRAIN | Miss Prop |
| --- | --- | --- | --- | --- | --- | --- |
| Group-LASSO  No missingness | 0.989 | 0.9889 | 3.96 | 3.98 | 50000 | 0 |
| Group-LASSO | 0.5329 | 0.7755 | 797.93 | 133.41 | 218 | 0.140 |
| MAGIC-LASSO (n trees = 3) | 0.5643 | 0.854 | 802.12 | 68.2 | 501 | 0.140 |
| MAGIC-LASSO (n trees = 4) | 0.5674 | 0.858 | 750.38 | 58.86 | 452 | 0.140 |
| MAGIC-LASSO (n trees = 5) | 0.5563 | 0.8446 | 747.29 | 66.02 | 324 | 0.140 |
| MAGIC-LASSO (n trees = 6) | 0.5537 | 0.8489 | 665.98 | 54.63 | 304 | 0.140 |
| MAGIC-LASSO (n trees = 7) | 0.5551 | 0.8509 | 681.15 | 54.38 | 336 | 0.140 |
| MAGIC-LASSO (n trees = 8) | 0.5528 | 0.8471 | 702.33 | 56.26 | 335 | 0.140 |
| MAGIC-LASSO (n trees = 9) | 0.557 | 0.8508 | 713.28 | 54.98 | 368 | 0.140 |
| MAGIC-LASSO (n trees = 10) | 0.5492 | 0.8417 | 726.22 | 67.02 | 363 | 0.140 |
| MAGIC-LASSO (n trees = 11) | 0.5515 | 0.8417 | 718.59 | 69.27 | 350 | 0.140 |
| MAGIC-LASSO (n trees = 12) | 0.5518 | 0.8422 | 703.66 | 65.79 | 359 | 0.140 |
| MAGIC-LASSO (n trees = 3) | 0.5609 | 0.6665 | 235.17 | 122.2 | 7371 | 0.141 |
| MAGIC-LASSO (n trees = 4) | 0.5245 | 0.6255 | 265.48 | 158.55 | 6956 | 0.141 |
| MAGIC-LASSO (n trees = 5) | 0.4907 | 0.5931 | 1068.82 | 908.74 | 7056 | 0.141 |
| MAGIC-LASSO (n trees = 6) | 0.4766 | 0.5821 | 528.08 | 391.55 | 7134 | 0.141 |
| MAGIC-LASSO (n trees = 7) | 0.4774 | 0.5865 | 282.52 | 170.33 | 7127 | 0.141 |
| MAGIC-LASSO (n trees = 8) | 0.4998 | 0.6117 | 269.43 | 163.21 | 7097 | 0.141 |
| MAGIC-LASSO (n trees = 9) | 0.4866 | 0.588 | 275.19 | 170.71 | 7367 | 0.141 |
| MAGIC-LASSO (n trees = 10) | 0.4316 | 0.5181 | 608.99 | 502.61 | 7365 | 0.141 |
| MAGIC-LASSO (n trees = 11) | 0.4353 | 0.5213 | 498.77 | 391.44 | 7389 | 0.141 |
| MAGIC-LASSO (n trees = 12) | 0.4372 | 0.5214 | 486.91 | 384.36 | 7398 | 0.141 |

Legend:

COR: correlation between observed and predicted outcomes for the missing outcomes

COR-All: correlation between observed and predicted outcomes for the full data set

MSE: mean squared error of prediction between observed and predicted outcomes for the missing outcomes

MSE-All: mean squared error of prediction between observed and predicted outcomes for the full dataset

N-Train: number of observations used to construct the model

Miss Prop: proportion of the total data set missing

n trees: number of tree utilized for cutting the clustering trees

*Shaded cells indicate scenarios with zero complete data cases

**Supplementary Table 7:** Simulation results for MAGIC-LASSO simulation, scenario 6.

| **Scenario 6** | COR | COR-All | MSE | MSE-all | N-TRAIN | Miss Prop |
| --- | --- | --- | --- | --- | --- | --- |
| Group-LASSO  No missingness | 0.9451 | 0.9448 | 3.97 | 3.99 | 50000 | 0 |
| Group-LASSO | 0.6316 | 0.7599 | 87.85 | 40.37 | 218 | 0.140 |
| MAGIC-LASSO (n trees = 3) | 0.7135 | 0.8627 | 33.01 | 16.22 | 910 | 0.140 |
| MAGIC-LASSO (n trees = 4) | 0.7217 | 0.8721 | 34.51 | 13.57 | 666 | 0.140 |
| MAGIC-LASSO (n trees = 5) | 0.6912 | 0.8403 | 40.83 | 19.16 | 363 | 0.140 |
| MAGIC-LASSO (n trees = 6) | 0.6886 | 0.8266 | 58.16 | 24.15 | 339 | 0.140 |
| MAGIC-LASSO (n trees = 7) | 0.6916 | 0.8294 | 49 | 22.36 | 421 | 0.140 |
| MAGIC-LASSO (n trees = 8) | 0.6993 | 0.8426 | 40.64 | 19.08 | 421 | 0.140 |
| MAGIC-LASSO (n trees = 9) | 0.7113 | 0.8575 | 31.19 | 14.68 | 491 | 0.140 |
| MAGIC-LASSO (n trees = 10) | 0.7063 | 0.8521 | 30.62 | 15.26 | 393 | 0.140 |
| MAGIC-LASSO (n trees = 11) | 0.7016 | 0.8467 | 38.85 | 17.78 | 372 | 0.140 |
| MAGIC-LASSO (n trees = 12) | 0.7016 | 0.847 | 38.63 | 17.73 | 369 | 0.140 |
| MAGIC-LASSO (n trees = 3) | 0.6653 | 0.6971 | 21.38 | 19.12 | 9328 | 0.141 |
| MAGIC-LASSO (n trees = 4) | 0.6449 | 0.679 | 26.34 | 23.98 | 8760 | 0.141 |
| MAGIC-LASSO (n trees = 5) | 0.6182 | 0.6529 | 32.76 | 29.46 | 9706 | 0.141 |
| MAGIC-LASSO (n trees = 6) | 0.6035 | 0.6393 | 37.07 | 33.51 | 9955 | 0.141 |
| MAGIC-LASSO (n trees = 7) | 0.5882 | 0.6224 | 42.61 | 39.9 | 9966 | 0.141 |
| MAGIC-LASSO (n trees = 8) | 0.6117 | 0.6417 | 27.56 | 25.92 | 9835 | 0.141 |
| MAGIC-LASSO (n trees = 9) | 0.5846 | 0.6144 | 32.67 | 30.82 | 10020 | 0.141 |
| MAGIC-LASSO (n trees = 10) | 0.5598 | 0.5906 | 29.59 | 27.66 | 11072 | 0.141 |
| MAGIC-LASSO (n trees = 11) | 0.5623 | 0.5938 | 28.19 | 26.22 | 11406 | 0.141 |
| MAGIC-LASSO (n trees = 12) | 0.5729 | 0.6021 | 57.3 | 52.88 | 10831 | 0.141 |

Legend:

COR: correlation between observed and predicted outcomes for the missing outcomes

COR-All: correlation between observed and predicted outcomes for the full data set

MSE: mean squared error of prediction between observed and predicted outcomes for the missing outcomes

MSE-All: mean squared error of prediction between observed and predicted outcomes for the full dataset

N-Train: number of observations used to construct the model

Miss Prop: proportion of the total data set missing

n trees: number of tree utilized for cutting the clustering trees

*Shaded cells indicate scenarios with zero complete data cases

**Supplementary Table 8:**

| **Supplementary Table 8:** Clustering details for rounds 1 (a) and 2 (b). For each iterative round, number of phenotypes, complete cases per cluster, and number of phenotype measures retained after applying the Group-Lasso to each cluster are shown.  **(a) Round 1** | | | | | |
| --- | --- | --- | --- | --- | --- |
| **Cluster** | **Number of Phenotypes** | **Number of Complete Cases** | **Phenotypes retained in AUDIT T** | **Phenotypes retained in AUDIT C** | **Phenotypes retained in AUDIT P** |
| 1 | 250 | 80839 | 28 | 24 | 52 |
| 2 | 12 | 19292 | 9 | 11 | 12 |
| 3 | 7 | 44941 | 7 | 7 | 7 |
| 4 | 6 | 16436 | 1 | 5 | 0 |
| 5 | 14 | 24888 | 9 | 8 | 8 |
| 6 | 2 | 27852 | 2 | 2 | 2 |
| 7 | 5 | 67766 | 5 | 5 | 5 |
| 8 | 5 | 0 | - | - | - |
| 9 | 11 | 225 | 5 | 6 | 5 |
| 10 | 31 | 45638 | 22 | 28 | 21 |
| 11 | 2 | 31086 | 2 | 1 | 2 |
| 12 | 9 | 120601 | 9 | 9 | 9 |

| **(b) Round 2** | | | |
| --- | --- | --- | --- |
| **AUDIT-T** |  |  |  |
| Cluster | Number of Phenotypes | Number of Complete Cases | Phenotypes Retained |
| 1 | 46 | 16363 | 21 |
| 2 | 17 | 4031 | 13 |
| 3 | 7 | 12680 | 7 |
| 4 | 5 | 36004 | 5 |
| 5 | 24 | 27737 | 19 |
| **AUDIT-C** |  |  |  |
| Cluster | Number of Phenotypes | Number of Complete Cases | Phenotypes Retained |
| 1 | 33 | 85210 | 17 |
| 2 | 19 | 5169 | 18 |
| 3 | 12 | 6312 | 8 |
| 4 | 7 | 12680 | 7 |
| 5 | 6 | 6608 | 5 |
| 6 | 29 | 33174 | 25 |
| **AUDIT-P** |  |  |  |
| Cluster | Number of Phenotypes | Number of Complete Cases | Phenotypes Retained |
| 1 | 69 | 16922 | 14 |
| 2 | 19 | 8244 | 10 |
| 3 | 7 | 12680 | 7 |
| 4 | 28 | 6612 | 23 |

**Supplementary Table 9:** Stepwise Group-Lasso application results for AUDIT-Total (a), AUDIT-Consumption (b), and AUDIT-Problems (c). For each step, the number of complete cases (subjects with complete data) in and the proportion of observations in the measured and unmeasured sets, as well as the phenotypic correlation for the prediction model built on that set, are shown. The highlighted line indicates the best performing model, with 30 phenotypes in AUDIT-Total, 18 in AUDIT-C, and 20 in AUDIT-P.

**(a) AUDIT-Total**

| **Number of Phenotypes** | **Number of Complete Cases** | **Number of Complete Cases (measured)** | **Number of Complete Cases (unmeasured)** | **Proportion of Measured Observations** | **Proportion of Unmeasured Observations** | **Ratio of Measured to Unmeasured** | **Phenotypic Correlation** |
| --- | --- | --- | --- | --- | --- | --- | --- |
| 1 | 502536 | 157162 | 345374 | 1 | 1 | 1 | NA |
| 2 | 502536 | 157162 | 345374 | 1 | 1 | 1 | 0.2684 |
| 3 | 501645 | 157089 | 344556 | 0.9995 | 0.9976 | 0.9981 | 0.3419 |
| 4 | 501639 | 157088 | 344551 | 0.9995 | 0.9976 | 0.9981 | 0.6044 |
| 5 | 501632 | 157088 | 344544 | 0.9995 | 0.9976 | 0.9981 | 0.6058 |
| 6 | 501515 | 157067 | 344448 | 0.9994 | 0.9973 | 0.9979 | 0.611 |
| 7 | 499725 | 156837 | 342888 | 0.9979 | 0.9928 | 0.9949 | 0.616 |
| 8 | 497760 | 156548 | 341212 | 0.9961 | 0.9879 | 0.9918 | 0.6081 |
| 9 | 489704 | 154505 | 335199 | 0.9831 | 0.9705 | 0.9872 | 0.6089 |
| 10 | 489654 | 154491 | 335163 | 0.983 | 0.9704 | 0.9872 | 0.6128 |
| 11 | 472443 | 149925 | 322518 | 0.954 | 0.9338 | 0.9789 | 0.613 |
| 12 | 453342 | 144398 | 308944 | 0.9188 | 0.8945 | 0.9736 | 0.6174 |
| 13 | 453339 | 144397 | 308942 | 0.9188 | 0.8945 | 0.9736 | 0.6187 |
| 14 | 433878 | 138101 | 295777 | 0.8787 | 0.8564 | 0.9746 | 0.6189 |
| 15 | 433378 | 137942 | 295436 | 0.8777 | 0.8554 | 0.9746 | 0.6197 |
| 16 | 432764 | 137767 | 294997 | 0.8766 | 0.8541 | 0.9744 | 0.6281 |
| 17 | 399391 | 131141 | 268250 | 0.8344 | 0.7767 | 0.9308 | 0.623 |
| 18 | 364427 | 125509 | 238918 | 0.7986 | 0.6918 | 0.8662 | 0.6035 |
| 19 | 331556 | 113971 | 217585 | 0.7252 | 0.63 | 0.8687 | 0.628 |
| 20 | 271230 | 95947 | 175283 | 0.6105 | 0.5075 | 0.8313 | 0.6155 |
| 21 | 215930 | 79510 | 136420 | 0.5059 | 0.395 | 0.7808 | 0.5544 |
| 22 | 163047 | 62004 | 101043 | 0.3945 | 0.2926 | 0.7416 | 0.5469 |
| 23 | 149450 | 56980 | 92470 | 0.3626 | 0.2677 | 0.7385 | 0.4953 |
| 24 | 149450 | 56980 | 92470 | 0.3626 | 0.2677 | 0.7385 | 0.5284 |
| 25 | 149450 | 56980 | 92470 | 0.3626 | 0.2677 | 0.7385 | 0.6179 |
| 26 | 149450 | 56980 | 92470 | 0.3626 | 0.2677 | 0.7385 | 0.6376 |
| 27 | 149450 | 56980 | 92470 | 0.3626 | 0.2677 | 0.7385 | 0.6376 |
| 28 | 91677 | 33330 | 58347 | 0.2121 | 0.1689 | 0.7966 | 0.6265 |
| 29 | 55671 | 19386 | 36285 | 0.1234 | 0.1051 | 0.8517 | 0.6366 |
| 30 | 26307 | 9862 | 16445 | 0.0628 | 0.0476 | 0.7588 | 0.639 |
| 31 | 26307 | 9862 | 16445 | 0.0628 | 0.0476 | 0.7588 | 0.637 |
| 32 | 26307 | 9862 | 16445 | 0.0628 | 0.0476 | 0.7588 | 0.637 |
| 33 | 26307 | 9862 | 16445 | 0.0628 | 0.0476 | 0.7588 | 0.637 |
| 34 | 12636 | 4904 | 7732 | 0.0312 | 0.0224 | 0.7175 | 0.6241 |
| 35 | 5844 | 2062 | 3782 | 0.0131 | 0.011 | 0.8346 | 0.6271 |
| 36 | 5844 | 2062 | 3782 | 0.0131 | 0.011 | 0.8346 | 0.6271 |
| 37 | 5844 | 2062 | 3782 | 0.0131 | 0.011 | 0.8346 | 0.6271 |
| 38 | 5844 | 2062 | 3782 | 0.0131 | 0.011 | 0.8346 | 0.6271 |
| 39 | 5844 | 2062 | 3782 | 0.0131 | 0.011 | 0.8346 | 0.6271 |
| 40 | 2612 | 852 | 1760 | 0.0054 | 0.0051 | 0.94 | 0.5728 |
| 41 | 0 | 0 | 0 | 0 | 0 | NA | NA |

**(b) AUDIT-Consumption**

| **Number of Phenotypes** | **Number of Complete Cases** | **Number of Complete Cases (measured)** | **Number of Complete Cases (unmeasured)** | **Proportion of Measured Observations** | **Proportion of Unmeasured Observations** | **Ratio of Measured to Unmeasured** | **Phenotypic Correlation** |
| --- | --- | --- | --- | --- | --- | --- | --- |
| 1 | 502536 | 157162 | 345374 | 1 | 1 | 1 | NA |
| 2 | 502536 | 157162 | 345374 | 1 | 1 | 1 | 0.2884 |
| 3 | 501645 | 157089 | 344556 | 0.9995 | 0.9976 | 0.9981 | 0.3471 |
| 4 | 501639 | 157088 | 344551 | 0.9995 | 0.9976 | 0.9981 | 0.6881 |
| 5 | 501522 | 157067 | 344455 | 0.9994 | 0.9973 | 0.9979 | 0.687 |
| 6 | 499529 | 156778 | 342751 | 0.9976 | 0.9924 | 0.9948 | 0.6881 |
| 7 | 496519 | 156224 | 340295 | 0.994 | 0.9853 | 0.9912 | 0.6922 |
| 8 | 488461 | 154161 | 334300 | 0.9809 | 0.9679 | 0.9868 | 0.6898 |
| 9 | 488411 | 154147 | 334264 | 0.9808 | 0.9678 | 0.9868 | 0.6901 |
| 10 | 475775 | 150955 | 324820 | 0.9605 | 0.9405 | 0.9792 | 0.6927 |
| 11 | 475775 | 150955 | 324820 | 0.9605 | 0.9405 | 0.9792 | 0.6935 |
| 12 | 461337 | 146357 | 314980 | 0.9312 | 0.912 | 0.9793 | 0.6896 |
| 13 | 461334 | 146356 | 314978 | 0.9312 | 0.912 | 0.9793 | 0.6932 |
| 14 | 442202 | 140174 | 302028 | 0.8919 | 0.8745 | 0.9805 | 0.6921 |
| 15 | 441698 | 140017 | 301681 | 0.8909 | 0.8735 | 0.9804 | 0.6979 |
| 16 | 441068 | 139835 | 301233 | 0.8898 | 0.8722 | 0.9803 | 0.6991 |
| 17 | 401599 | 127090 | 274509 | 0.8087 | 0.7948 | 0.9829 | 0.7018 |
| 18 | 332697 | 107054 | 225643 | 0.6812 | 0.6533 | 0.9591 | 0.7053 |
| 19 | 267747 | 89546 | 178201 | 0.5698 | 0.516 | 0.9056 | 0.6976 |
| 20 | 229101 | 77294 | 151807 | 0.4918 | 0.4395 | 0.8937 | 0.6938 |
| 21 | 181950 | 64355 | 117595 | 0.4095 | 0.3405 | 0.8315 | 0.6074 |
| 22 | 147982 | 53641 | 94341 | 0.3413 | 0.2732 | 0.8003 | 0.6106 |
| 23 | 109901 | 38096 | 71805 | 0.2424 | 0.2079 | 0.8577 | 0.5998 |
| 24 | 84491 | 28471 | 56020 | 0.1812 | 0.1622 | 0.8954 | 0.6134 |
| 25 | 67376 | 22807 | 44569 | 0.1451 | 0.129 | 0.8892 | 0.6026 |
| 26 | 50148 | 17732 | 32416 | 0.1128 | 0.0939 | 0.8319 | 0.6023 |
| 27 | 50148 | 17732 | 32416 | 0.1128 | 0.0939 | 0.8319 | 0.6021 |
| 28 | 46032 | 16357 | 29675 | 0.1041 | 0.0859 | 0.8256 | 0.5732 |
| 29 | 46032 | 16357 | 29675 | 0.1041 | 0.0859 | 0.8256 | 0.5975 |
| 30 | 46032 | 16357 | 29675 | 0.1041 | 0.0859 | 0.8256 | 0.6477 |
| 31 | 46032 | 16357 | 29675 | 0.1041 | 0.0859 | 0.8256 | 0.6609 |
| 32 | 46032 | 16357 | 29675 | 0.1041 | 0.0859 | 0.8256 | 0.6613 |
| 33 | 30054 | 8731 | 21323 | 0.0556 | 0.0617 | 1.1113 | 0.6677 |
| 34 | 25683 | 7353 | 18330 | 0.0468 | 0.0531 | 1.1344 | 0.6628 |
| 35 | 14602 | 4503 | 10099 | 0.0287 | 0.0292 | 1.0206 | 0.6663 |
| 36 | 9876 | 2979 | 6897 | 0.019 | 0.02 | 1.0535 | 0.6643 |
| 37 | 7380 | 2215 | 5165 | 0.0141 | 0.015 | 1.0611 | 0.6626 |
| 38 | 7380 | 2215 | 5165 | 0.0141 | 0.015 | 1.0611 | 0.6632 |
| 39 | 7380 | 2215 | 5165 | 0.0141 | 0.015 | 1.0611 | 0.663 |
| 40 | 7380 | 2215 | 5165 | 0.0141 | 0.015 | 1.0611 | 0.663 |
| 41 | 3311 | 1066 | 2245 | 0.0068 | 0.0065 | 0.9583 | 0.5482 |
| 42 | 1586 | 574 | 1012 | 0.0037 | 0.0029 | 0.8023 | 0.5793 |
| 43 | 760 | 256 | 504 | 0.0016 | 0.0015 | 0.8959 | 0.6511 |
| 44 | 0 | 0 | 0 | 0 | 0 | NA | NA |

**(c) AUDIT-Problems**

| **Number of Phenotypes** | **Number of Complete Cases** | **Number of Complete Cases (measured)** | **Number of Complete Cases (unmeasured)** | **Proportion of Measured Observations** | **Proportion of Unmeasured Observations** | **Ratio of Measured to Unmeasured** | **Phenotypic Correlation** |
| --- | --- | --- | --- | --- | --- | --- | --- |
| 1 | 502536 | 157162 | 345374 | 1 | 1 | 1 | NA |
| 2 | 501645 | 157089 | 344556 | 0.9995 | 0.9976 | 0.9981 | 0.2076 |
| 3 | 501639 | 157088 | 344551 | 0.9995 | 0.9976 | 0.9981 | 0.3199 |
| 4 | 501629 | 157088 | 344541 | 0.9995 | 0.9976 | 0.9981 | 0.3301 |
| 5 | 501512 | 157067 | 344445 | 0.9994 | 0.9973 | 0.9979 | 0.3386 |
| 6 | 499724 | 156837 | 342887 | 0.9979 | 0.9928 | 0.9949 | 0.3548 |
| 7 | 497759 | 156548 | 341211 | 0.9961 | 0.9879 | 0.9918 | 0.3438 |
| 8 | 489703 | 154505 | 335198 | 0.9831 | 0.9705 | 0.9872 | 0.3494 |
| 9 | 467267 | 148214 | 319053 | 0.9431 | 0.9238 | 0.9796 | 0.359 |
| 10 | 446935 | 141689 | 305246 | 0.9015 | 0.8838 | 0.9803 | 0.3706 |
| 11 | 446291 | 141504 | 304787 | 0.9004 | 0.8825 | 0.9801 | 0.3655 |
| 12 | 444751 | 141046 | 303705 | 0.8975 | 0.8794 | 0.9798 | 0.3718 |
| 13 | 357287 | 117944 | 239343 | 0.7505 | 0.693 | 0.9234 | 0.3797 |
| 14 | 280039 | 97110 | 182929 | 0.6179 | 0.5297 | 0.8572 | 0.3866 |
| 15 | 252356 | 88187 | 164169 | 0.5611 | 0.4753 | 0.8471 | 0.4084 |
| 16 | 252356 | 88187 | 164169 | 0.5611 | 0.4753 | 0.8471 | 0.4376 |
| 17 | 252356 | 88187 | 164169 | 0.5611 | 0.4753 | 0.8471 | 0.4569 |
| 18 | 252356 | 88187 | 164169 | 0.5611 | 0.4753 | 0.8471 | 0.4674 |
| 19 | 252356 | 88187 | 164169 | 0.5611 | 0.4753 | 0.8471 | 0.4685 |
| 20 | 147063 | 48890 | 98173 | 0.3111 | 0.2843 | 0.9138 | 0.4785 |
| 21 | 71542 | 26002 | 45540 | 0.1654 | 0.1319 | 0.797 | 0.4623 |
| 22 | 71542 | 26002 | 45540 | 0.1654 | 0.1319 | 0.797 | 0.4623 |
| 23 | 71542 | 26002 | 45540 | 0.1654 | 0.1319 | 0.797 | 0.4623 |
| 24 | 71542 | 26002 | 45540 | 0.1654 | 0.1319 | 0.797 | 0.4623 |
| 25 | 39805 | 13484 | 26321 | 0.0858 | 0.0762 | 0.8883 | 0.4663 |
| 26 | 39805 | 13484 | 26321 | 0.0858 | 0.0762 | 0.8883 | 0.4663 |
| 27 | 39805 | 13484 | 26321 | 0.0858 | 0.0762 | 0.8883 | 0.4663 |
| 28 | 39805 | 13484 | 26321 | 0.0858 | 0.0762 | 0.8883 | 0.4663 |
| 29 | 39805 | 13484 | 26321 | 0.0858 | 0.0762 | 0.8883 | 0.4663 |
| 30 | 17562 | 5429 | 12133 | 0.0345 | 0.0351 | 1.017 | 0.3982 |
| 31 | 0 | 0 | 0 | 0 | 0 | NA | NA |

**Supplementary Table 10:**

| **Supplementary Table 10:** GCTA estimated heritabilities ($h^{2}$) and genetic correlations ($r_{g}$) for AUDIT-Total. $r_{g}$ calculated as the genetic correlation between the observed and predicted scores within the measured set only. | | | | | | | |
| --- | --- | --- | --- | --- | --- | --- | --- |
| **Set** | **Outcome** | **Sex** | $h^{2}$ | **SE (**$h^{2}$**)** | **N** | $r_{g}$ | **SE (**$r_{g}$**)** |
| Measured | Observed | Male | 0.139 | 0.0108 | 50,912 | 0.863 | 0.040 |
| Measured | Predicted | Male | 0.091 | 0.0105 | 50,912 |  |  |
| Measured | Observed | Female | 0.109 | 0.0087 | 64,768 | 0.884 | 0.032 |
| Measured | Predicted | Female | 0.092 | 0.0084 | 64,768 |  |  |
| Unmeasured | Predicted | Male | 0.089 | 0.0052 | 109,916 |  |  |
| Unmeasured | Predicted | Female | 0.108 | 0.0048 | 122,159 |  |  |

Legend:

$h^{2}$: heritability

$r_{g}$: genetic correlation

SE: standard error

**Supplementary Figure 1**

**Supplementary Figure 1:** Illustration of clustering and Group-LASSO procedures within the MAGIC-LASSO. The hierarchical clustering steps divides the predictor space into non-overlapping subsets based on missingness. The Group-LASSO is applied to each cluster and variables retained by the algorithm in each set are aggregated.

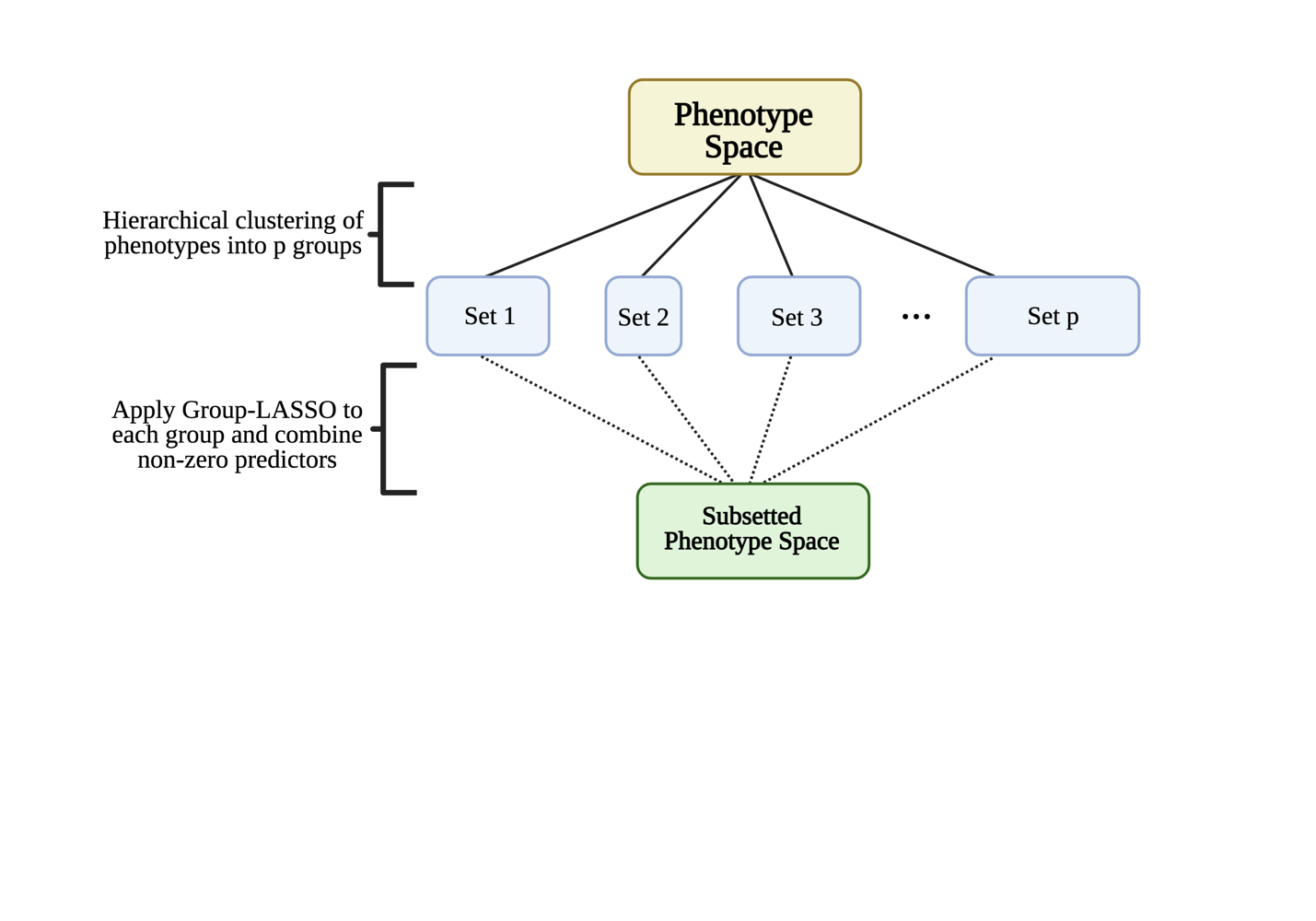

**Supplementary Figure 2:** Flowchart of filtering in the UKB application set. The number of variables remaining after each filtering metric is applied is shown for the UKB data application example.

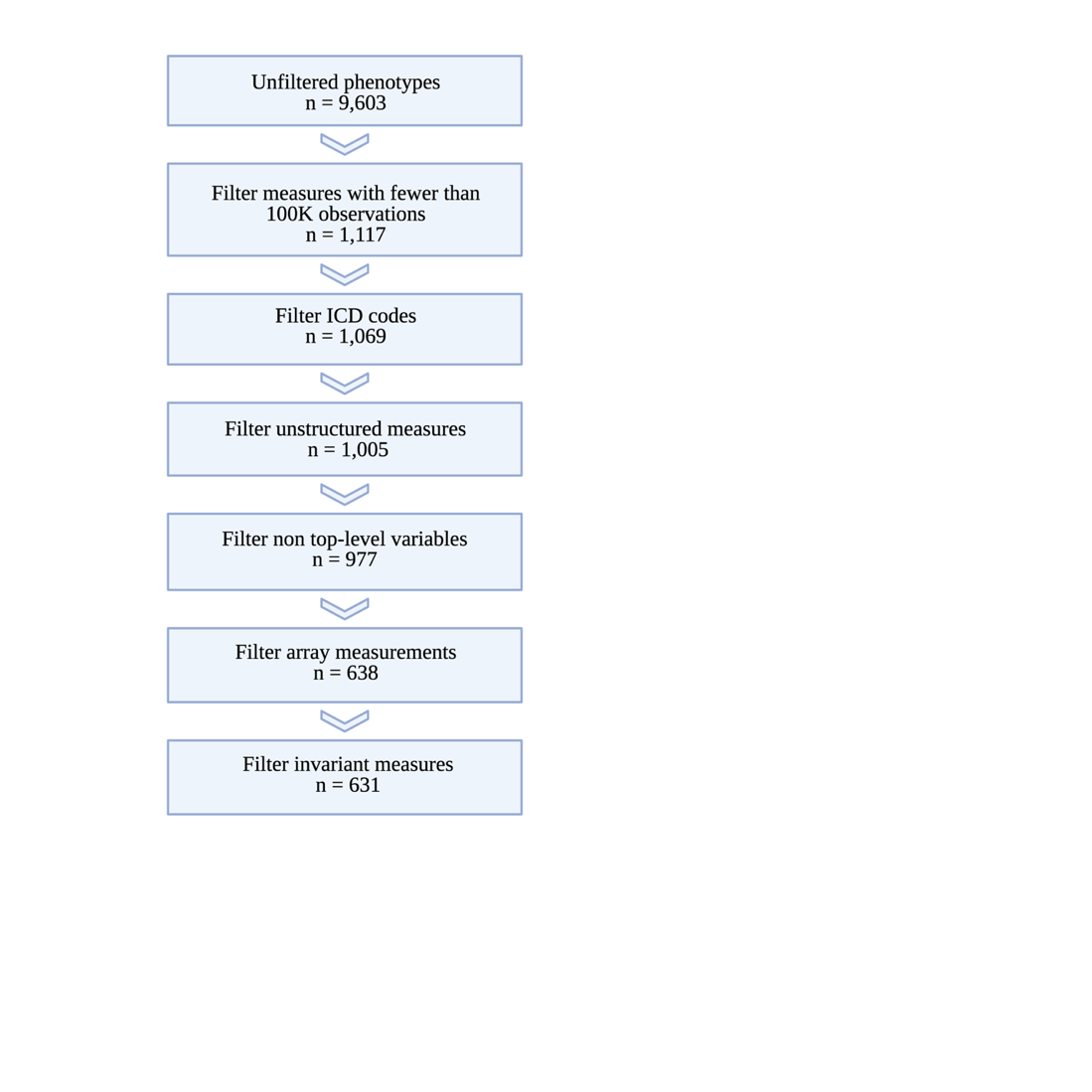

**Supplementary Figure 3:** Stepwise model results. Phenotypic correlations and ratio of the proportion of total measured to unmeasured observations in the model set for (a) AUDIT-Total, (b) AUDIT-Consumption, and (c) AUDIT-Problems. Final selected model highlighted in blue.

(a) AUDIT-Total

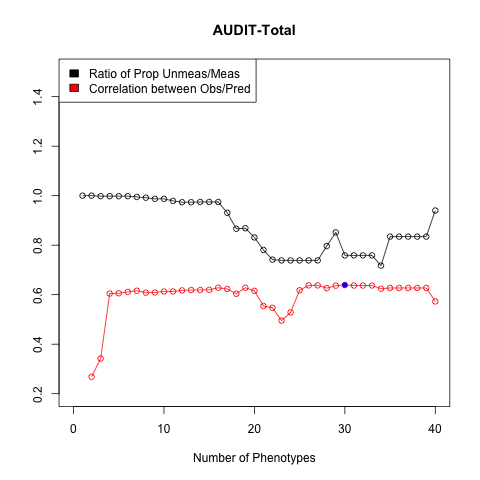

(b) AUDIT-Consumption

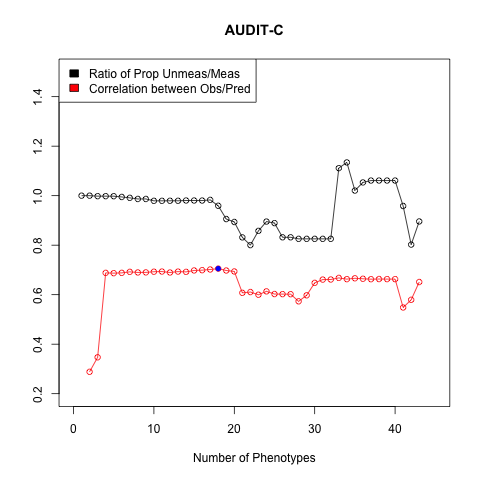

(c) AUDIT-Problems

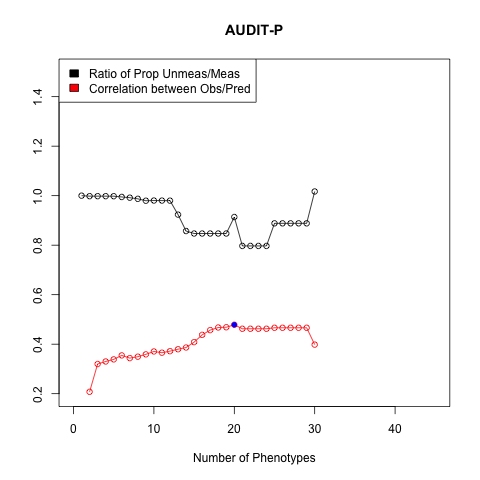
